# Towards an optimal monoclonal antibody with higher binding affinity to the receptor-binding domain of SARS-CoV-2 spike proteins from different variants

**DOI:** 10.1101/2022.01.04.474958

**Authors:** Andrei Neamtu, Francesca Mocci, Aatto Laaksonen, Fernando L. Barroso da Silva

**Affiliations:** Department of Physiology, “Grigore T. Popa” University of Medicine and Pharmacy of Iasi, Str. Universitatii nr. 16, 700051 Iasi, România; Centre of Advanced Research in Bionanoconjugates and Biopolymers, PetruPoni Institute of Macromolecular Chemistry AleeaGrigoreGhica-Voda, 41A, 700487 Iasi, Romania; University of Cagliari, Department of Chemical and Geological Sciences, Campus Monserrato, SS 554 bivio per Sestu, 09042 Monserrato, Italy; Department of Materials and Environmental Chemistry, Arrhenius Laboratory, Stockholm University, SE-106 91 Stockholm, Sweden; State Key Laboratory of Materials-Oriented and Chemical Engineering, Nanjing Tech University, Nanjing, 210009, P. R. China; Department of Engineering Sciences and Mathematics, Division of Energy Science, Luleå University of Technology, SE-97187 Luleå, Sweden; Universidade de São Paulo, Departamento de CiênciasBiomoleculares, Faculdade de CiênciasFarmacêuticas de Ribeirão Preto, Av. café, s/no – campus da USP, BR-14040-903 – Ribeirão Preto – SP, Brazil; Department of Chemical and Biomolecular Engineering, North Carolina State University, Raleigh,North Carolina 27695, United States

**Author notes:** Corresponding author ( and/or).

## Abstract

A highly efficient and robust multiple scales *in silico* protocol, consisting of atomistic constant charge Molecular Dynamics (MD), constant-charge coarse-grain (CG) MD and constant-pH CG Monte Carlo (MC), has been used to study the binding affinities, the free energy of complexation of selected antigen-binding fragments of the monoclonal antibody (mAbs) CR3022 (originally derived from SARS-CoV-1 patients almost two decades ago) and 11 SARS-CoV-2 variants including the wild type. CR3022 binds strongly to the receptor-binding domain (RBD) of SARS-CoV-2 spike protein, but chooses a different site rather than the receptor-binding motif (RBM) of RBD, allowing its combined use with other mAbs against new emerging virus variants. Totally 235,000 mAbs structures were generated using the RosettaAntibodyDesign software, resulting in top 10 scored CR3022-RBD complexes with critical mutations and compared to the native one, all having the potential to block virus-host cell interaction. Of these 10 finalists, two candidates were further identified in the CG simulations to be clearly best against all virus variants, and surprisingly, all 10 candidates and the native CR3022 did exhibit a higher affinity for the Omicron variant with its highest number of mutations (15) of them all considered in this study. The multiscale protocol gives us a powerful rational tool to design efficient mAbs. The electrostatic interactions play a crucial role and appear to be controlling the affinity and complex building. Clearly, mAbs carrying a lower net charge show a higher affinity. Structural determinants could be identified in atomistic simulations and their roles are discussed in detail to further hint at a strategy towards designing the best RBD binder. Although the SARS-CoV-2 was specifically targeted in this work, our approach is generally suitable for many diseases and viral and bacterial pathogens, leukemia, cancer, multiple sclerosis, rheumatoid, arthritis, lupus, and more.

## INTRODUCTION

World has been in the grip of the “Covid-19” pandemic, caused by the severe acute respiratory syndrome coronavirus 2 (SARS-CoV-2), for roughly two years with no corner on Earth being saved. To date, more than 281 million cases and 5.4 million deaths are reported with no end in sight [https://www.who.int/] at the time of writing. The pandemic has quickly mobilized the scientific communities, pharmacological companies, and governmental health agencies. Vaccines have been developed in the shortest possible time and manufactured in large quantities on a global scale (Mallapaty et al., 2021). Yet, the number of people to be vaccinated remains quite large. How the pandemics will develop largely depends on the populations becoming vaccinated worldwide in combination with non-pharmacological measures. Unfortunately, on one hand, there is strong resistance to accepting the vaccine, and on the other hand, poor countries do not have supplies for their populations. This worsens the situation as it gives time for new mutations of the virus to develop with a relatively high potential for breakthrough cases (Gupta and Topol, 2021). Consequently, it requires continuously new research and constantly new treatment strategies. To be able to combat the infections, and hopefully to finally put an end to the pandemics, it is important to investigate the genomics, molecular structure and dynamics, mechanisms of binding, and the life cycle of the viruses.

Since the discovery of Coronaviruses 90 years ago(V’kovski et al., 2021), more than 30 different coronaviruses have been discovered so far, 6 of which infect humans. During this millennium, three zoonotic outbreaks of severe acute respiratory syndrome coronaviruses have taken place: SARS-CoV in 2002-2003, Middle East respiratory syndrome coronavirus (MERS-CoV) in 2012, and the one we are currently experiencing, SARS-CoV-2, which generated the most critical pandemic experienced so far by humans(Fung et al., 2021).

Coronaviruses are round or elliptic particles around 120 nm in diameter that can easily change their form. They consist of positive-sense single-stranded +ssRNA with 3’ and 5’ terminus cap, extending up to ca. 30 kilobases coated by two types of nucleocapsid proteins and embedded by a double-layer phospholipid membrane. The surface of coronaviruses is coated by glycoproteins. There are four types of glycoproteins all taking part in the pathogenesis: spike (S), membrane (M), nucleocapsid (N), and envelope (E). The S-protein is a homotrimer sticking out of the viral surface. There is a specific site for the protease furin to cleavage it in the spike protein(Whittaker, 2021), and an open reading frame1a/b differentiating it from other RNA viruses. The furin site is particularly mutation-prone (V’kovski et al., 2021). The spike protein is highly flexible, being even able to rotate and swing, and compared to other coronaviruses, where the spike protein is rather rigid, it is much more efficient in binding. In the top of spike protein, there is the RBD which binds to the angiotensin-converting enzyme 2 (ACE2) receptor of the host cell to promote an entry to infect the human cell assisted with host factors such as the cell surface serine protease TMPRSS2 or cathepsin L by fusing with the cellular or endosomal membrane (Li, 2016; Li et al., 2005; V’kovski et al., 2021; Walls et al., 2020)]. Each chain of the S protein contains two subunits, S1 and S2. They become cleaved by TMPRSS2 both having important roles in the fusion to the host cell. S1 subunit leaks out from the S2unit facilitating the membrane fusion between viral and cell membranes by hydrophobic amino acids which become buried in the cell membrane of the host cell. Some studies show that SARS-CoV-2 has a much higher affinity and exhibits stronger interaction for ACE2 than the RBD of the previous SARS-CoV-1 (Tian et al., 2020). Other works measured similar affinities (Brielle et al., 2020). Nevertheless, the RBD region of the S protein is obviously an ideal target for vaccines to induce host immune response and generate neutralizing antibodies (Mallapaty et al., 2021; Prates-Syed et al., 2021). New variants tend to appear in the S1 subunit, for example, the Delta variant has several mutations there and also some specifically in RBD (Khateeb et al., 2021). The Alpha variant has ten changes in the spike protein sequence making the RBD stay in a position that helps the virus to enter the cell. The trimeric spike has several conformations (open and closed) and orientations before fusion, and once bound it undergoes a structural transition forming a narrow bridge which also protects the virus from the immune defense (Ismail and Elfiky, 2020). The probability to be in these two different conformational states (open and closed) is strongly dependent on the variants as it was seen in experimental and theoretical works (Giron et al., 2021; Moreira et al., 2020; Yuan et al., 2020). There seems to be a correlation between this probability and the tendency of either asymptomatic or symptomatic clinical cases (Chun Huai Luo et al., 2021; Giron et al., 2021).

While inside the cell, the virus can still be trappedby antiviral proteins but if not, the RNA is uncoated, released, and translated by two large open frames ORF1a and ORF1b resulting in polyproteins pp1a and pp1b, thereafter processed to non-structural proteins (nsps) to prepare the viral replication and transcription complex. First takes place the biogenesis of double-membrane vesicles (DMV) and convoluted membranes (CM) as well as small double-membrane spherules based on the expression of the nsps to provide a protective environment for the replication and transcription of the RNA sub-genome. The entire positive +ssRNA is also transcribed producing full-length negative -ssRNA, serving as a template to replicate the viral genome. The translated structural proteins (S, E, M, N) are enclosed in endoplasmic reticulum (ER) membranes and moved to the Golgi intermediate compartment (ERGIC) where the genomic RNA gets again encapsulated by N proteins. Finally, the viruses are spit out from the infected cell by exocytosis and are ready to infect new host cells (V’kovski et al., 2021). Many details are still not completely known concerning the virus inside the cell.

There are several therapy strategies that in principle can be used to combat SARS-CoV-2 infections. For example, in developing the vaccines, the main choices space from mRNA vaccines releasing a synthetic mRNA sequence encoding the virus spike protein from a nano-particle to virus vector vaccines where DNA encoding the spike protein is embedded in a viral vector to enter the cell (Mallapaty et al., 2021; Prates-Syed et al., 2021).

Despite some promising medicines (Reis et al., 2022) including PAXLOVID, there is still no effective drug against Covid-19 but neutralizing monoclonal antibodies (mAbs) are currently the most attractive alternative. One approach that has been used successfully in the past to treat various infectious diseases is convalescent plasma therapy (CPT) (Keller and Stiehm, 2000). This treatment uses plasma collected from COVID-19 recovering patients (those that have been cured successfully) to be administered to recipient patients that still did not develop a protective immune response. Neutralizing antibodies (Abs) from the donor’s plasma helps in reducing the viral load in the recipient. Apart from neutralization, other mechanisms may also be responsible for the protective effect of CPT. Some studies revealed the presence in the plasma of COVID-19 convalescent patients of Abs capable of inducing antibody-dependent cell cytotoxicity (ADCC) (Tso et al., 2021), phagocytosis (Natarajan et al., 2021), and complement activation (Natarajan et al., 2021). Despite its success, one important limitation of CPT in the treatment of COVID-19 is represented by the diversity of virus variants found in the population which makes the selection of donors difficult. The large variation of Abs level in different plasma samples and serum incompatibility of recipients are yet another issue that has to be carefully evaluated when using CPT (Li et al., 2020; Salazar et al., 2020). On the other hand, mAbs or polyclonal antibody cocktails can be specifically engineered and biotechnologically produced to fight SARS-CoV-2. Spike protein is the primary target of these mAbs with four classes of them being described to date depending on the location of their target epitope on the S protein (Kumar et al., 2021). Regardless of their source, either from convalescent blood or industrially produced, human mAbs are safe therapeutic tools and can be produced quickly. There are already more than 50 commercially available mAbs approved for the treatment of other inflammatory and immune disorders and other infectious pathogens. Several candidates against SARS-CoV-2 are by now in different trial phases (Chen et al., 2021; Hastie et al., 2021). Patents have already been deposited (e.g, US 2021/0292393), and some mAbs are approved by regulatory agencies such as the FDA (Boggiano et al., 2021; Rubin, 2021).

Among available mAbs, CR3022 is a class IV mAb that does not bind to an RBD epitope that overlaps the ACE2 binding site, but to a conserved region in RBD when the spike homotrimer is in “up” (or open) configuration exposing the RBD to interact with either ACE2 or binder molecules (Barnes et al., 2020; Giron et al., 2021; Yan et al., 2021). Exploiting such conserved regions in RBD (despite requiring the spike protein to be at the up/open state), class IV mAbs have thus broad neutralizing activity against SARS-CoV-2, its variants, and other related coronaviruses (e.g. SARS-CoV) (Barnes et al., 2020). The broad sarbecovirus neutralizing activity from CR3022 has already been confirmed even for the Omicron variant (Cao et al., 2021). Due to the abrupt appearance of new variants of concerns (VOCs) in a short time, improving a mAb that has its epitope on a conserved region of the RBD may constitute an advantage for developing biopharmaceuticals broadly effective against different variants of the virus. Moreover, attention has recently been turned again to the CR3022 antibody as a promising candidate for COVID-19 treatment and prevention as new experimental and computational evidence became available. The experimental study of Tian et al. (Tian et al., 2020) and the computational study of Nguyen et al. (Nguyen et al., 2021) showed that CR3022 binds RBD of SARS-CoV-2 with high affinity (K_D_ ∼ 6.3 to 3 nM) in contrast with the results of Yuan et al.(Yuan et al., 2020) which gave a much lower affinity of binding (K_D_ ∼ 115 nM). Other theoretical works showed similar binding affinities with a tendency for stronger complexes formed with RBD SARS-CoV-1 (Corrêa Giron et al., 2020). Despite these inconsistencies regarding the affinity of CR3022 for SARS-CoV-2 S-protein, this mAb is nevertheless an appealing candidate to be subjected to affinity maturation, especially due to its cross-reactivity. Therefore, the CR3022 antibody has been chosen in this study and will be the central part of our discussions.

Traditionally, mAbs production is based on immunization by hybridoma technology which uses an optimized immunogenic specific antigen to infect a host animal and then to isolate its short-lived mature B-cells from the spleen which can produce antigen-specific mAbs. These specific B-cells are then fused with immortal myeloma cells to obtain hybridomas which are able to secrete large amounts of the specific mAb (Köhler and Milstein, 1975; Zaroff and Tan, 2019). Promisingly, mAbs discovery for SARS-CoV-2 has benefited more recently also from *in vitro* approaches like phage (Noy-Porat et al., 2020) and yeast display (Cortese and Neufeldt, 2021) technologies. Despite their tremendous positive impact on therapeutic mAbs development, all these techniques present also challenges and limitations whose description falls beyond the scope of this paper but can be found in two excellent recent reviews (Laustsen et al., 2021; Parray et al., 2020). Both hybridoma and *in vitro* techniques sometimes lead to mAbs with lower than expected affinity and specificity (Foote and Eisen, 1995; González-Fernández et al., 2020). To alleviate this drawback, supplementary *in vitro* affinity maturation is used to increase the potency of the designed antibodies resulting in lower injected doses and side effects (Lim et al., 2016; Persson et al., 2018). The two approaches in use today in this respect include random mutagenesis (Tachioka et al., 2016) and chain shuffling/site-directed mutagenesis (Lou and Marks, 2010). Both of them introduce mutations in the complementarity-determining regions (CDR) sequences (the part of the variable chains in immunoglobulins), either at random or at specific positions, followed by affinity screening of the expressed mutant using display technologies to select the best-improved antibody (Kim et al., 2014). Although there are successful studies that have used *in vitro* affinity maturation alone, several challenges related to display technologies may limit its efficient applicability. These include the limited sub-space of the possible mutations effectively accessible to display approaches (Tiller et al., 2017), the time consuming and laborious process of building sub-libraries, and the risk of non-specific binding or loss of stability (Julian et al., 2017) and not in the last place the long time needed and prohibitively expensive price to complete experiments.

To successfully develop potent therapeutic Abs many recent studies employ, alongside experimental *in vitro* techniques, computational approaches which can provide detailed structural information on atomic-scale regarding the antibody-epitope interactions and to predict possible improvements (Lippow et al., 2007). The large number of such studies emphasizes that computational tools become routinely used for complementing experiments in the Ab design process. At present, a large diversity of algorithms for computational Ab studies are available, from those that predict the structure of an Ab based on its primary sequence only (i.e. antibody modeling) (Weitzner et al., 2017), to the ones that make use of detailed structural information from X-ray diffraction data to design improved antibodies from existent ones (i.e. antibody design) (Chowdhury et al., 2020). Availability of experimental structural data regarding the complex between the Ab to be improved and its antigen (Ag) greatly improves the prediction accuracy although antibody-antigen computationally modeled structures could be used as well (Sivasubramanian et al., 2009). In short, the general workflow of computational design of novel paratopes includes: (i) generation of new CDRs from experimental antibody databases and their grafting/modeling on the antibody framework, (ii) CDRs sequence redesign introducing mutations, (iii) antibody-antigen docking according to the known epitope and relative orientation of the partners, (iv) binding energy evaluation using a scoring function. Eventually, these steps are performed in many rounds to encounter a better binding antibody. The four generally used algorithms available today include: RosettaAntibodyDesign (Adolf-Bryfogle et al., 2018), AbDesign (Lapidoth et al., 2015), OptCDR (Pantazes and Maranas, 2010) and OptMAVEn (Chowdhury et al., 2018). In addition to this general protocol, other molecular modeling techniques can be applied to improve the accuracy of predictions. Molecular dynamics (MD) simulations, with more than 60 years of history (Alder and Wainwright, 1959; Rahman, 1964), represent a well-established method for analyzing the physical movement of atoms and molecules, with a long history of applicability in biological sciences (Barroso da Silva et al., 2020; Hollingsworth and Dror, 2018; Karplus and McCammon, 2002; McCammon et al., 1977). It has the advantage of introducing the time dimension in the simulations, this way being able to capture a wide variety of biological processes like conformational changes or protein folding and how biological molecules will respond to post-translational modifications or mutations. Atomistic MD has been successfully used in the past in Ab design studies for affinity maturation of camelid nanobodies against alpha-synuclein, a weak immunogenic antigen (Mahajan et al., 2018), of bevacizumab antibody for an increased affinity to vascular endothelial growth factor A (VEGF-A) (Corrada and Colombo, 2013), and of a toll-like receptor (TLR4) targeting Ab (Ahmad et al., 2021), to mention just a few examples. However, different steps of Ab-Ag interaction take place on different time scales, which cannot be effectively covered by conventional atomistic simulations only.

In this paper, we have used a *multiscale* approach for *in silico* affinity maturation of the mAb CR3022 against RBD of SARS-CoV-2 in order not only to improve binding but also to understand the molecular determinants on different time/length scales which can be exploited to further modify and modulate the binding affinity. Our approach included an initial conventional Ab design stage followed by complementary evaluations of the best candidates using constant-pH Monte Carlo (MC) calculations, constant-charge coarse-grain (CG) MD calculations, and constant-charge atomistic MD simulations. We then selected, as the result of the affinity maturation process, the candidate that showed improved affinity in all four types of evaluations. One strong point in favor of our approach is that it uses complementary methods for binding affinity estimation, with different physical bases ranging from rigid-body long-range interaction assessment to local conformational rearrangements upon intimate Ab-Ag binding. Furthermore, our multiscale approach allowed us to test our candidates against 11 different strains of the SARS-CoV-2 virus. We identified key residues important for binding located either at the interaction interface or distant to it, together with mutations that improve binding like S35K (L1), S72E (L1), Y110R (L3), and Y39W (H1). Also, the total net charge of the mAb proved to be an important parameter that should be considered when aiming to design better binders for RBD of SARS-CoV-2.

## THEORETICAL METHODS

Modern virology, immunology, and related fields are expanding their achievements incorporating theoretical approaches in their daily practice (Ibrahim et al., 2018; Sato et al., 2013; Sharma et al., 2015). In fact, computer simulations have been playing a key role in many biological systems including pharma and viruses (Barroso da Silva et al., 2020; Germain et al., 2011; Schlick and Portillo-Ledesma, 2021). Examples are found in several study cases from the understanding of capsid formation to the design of antiviral molecules (Boulard and Bressanelli, 2021; Francés-Monerris et al., 2020; Keretsu et al., 2020; Mondal and Warshel, 2020; Poveda-Cuevas et al., 2021, 2020; Rapaport, 2018). Canonical simulation methods well established in more fundamental scientific areas and material sciences are now routinely used to enhance the understanding of the biomolecular interactions involved in infectious diseases. Following a more biophysical approach, classical MD and MC methods are techniques that have a long history of successful applications in many scientific problems as standalone theoretical studies or in combination with experimental techniques (Barroso da Silva et al., 2020; Dascalu et al., 2017; Engelbrecht et al., 2022; Gunsteren et al., 1994; Lyubartsev and Laaksonen, 2021; Minea et al., 2016; Steinhauser and Hiermaier, 2009; van Gunsteren and Dolenc, 2012). Other bioinformatics algorithms are completing theoretical resources and helping to address challenging problems in the biomolecular world (Andreatta and Nielsen, 2018; Ibrahim et al., 2018; Pappas et al., 2021; Sironi and Kaderali, 2021).

### Some previous theoretical studies with CR3022

As mentioned above, CR3022 has been suggested as a promising therapeutic option to neutralize SARS-CoV-2 (Barnes et al., 2020; Tian et al., 2020; Wu et al., 2020; Yuan et al., 2020), and is often explored in theoretical studies (Corrêa Giron et al., 2020; Ding et al., 2021; Nguyen et al., 2021; Shariatifar and Farasat, 2021). In a pioneer computational study at the very beginning of the pandemic, using constant-pH MC simulations, Giron, Laaksonen, and Barroso da Silva showed that CR3022 known to bind to SARS-CoV-1 RBD could also bind to SARS-CoV-2 RBD. They also mapped the epitopes and identified the importance of electrostatic interactions for this Ab-Ag interface (Corrêa Giron et al., 2020). Following different routes, Ding et al. (Ding et al., 2021) have proposed an efficient and reliable computational screening method based on “Molecular Mechanics Poisson-Boltzmann” surface area to evaluate the binding free energy between SARS-CoV-2 RBD and ACE2 together with CR3022 and CB6 in good agreement with reported experimental values. Their scheme identifies the key residues to increase hydrophobicity and also that changing the sign of charged residues from positive to negative can increase the binding affinity. In comparison with standard MM/PBSA, their method is more accurate due to the introduction of electrostatic energy in the scheme.

Lagoumintzis et al. (Lagoumintzis et al., 2021) used *in silico* methods in their studies of the recent hypothesis that SARS-CoV2 would interact with nicotinic acetylcholine receptors (nAChRs) and disrupt the regulation of nicotinic cholinergic system (NCS) and the cholinergic anti-inflammatory pathway. They used ROSETTA and multi-template homology modeling to a sequence from a snake venom toxin for structure prediction of the extracellular domains of nAChRs (“toxin binding site”). Using the “High Ambiguity Driven protein-protein DOCKing” (HADDOCK) approach (van Zundert et al., 2016), they found a protective role of nicotine and other cholinergic agonists and observed CR3022 and other similar mAbs showing an increased affinity for SARS-CoV-2 Spike glycoprotein. To study the molecular mechanisms in SARS-CoV-2 S protein binding with several Abs, Verkhivker and Di Paola (Verkhivker and Di Paola, 2021) performed all-atom and CG simulations with mutational sensitivity mapping using the BeAtMuSiC approach and perturbation response scanning (PRS) profiling of SARS-CoV-2 receptor-binding domain complexed with CR3022 and CB6 antibodies, complementing it with a network modeling analysis of the residue interactions. Their results provide insight into allosteric regulatory mechanisms of SARS-CoV-2 S proteins, where the Abs are modulating the signal communication. This provides a strategy to target specific regions of allosteric interactions therapeutically. Recently, Riahi et al. (Riahi et al., 2021) presented a combined physics-based and machine learned-based computational Ab engineering platform to improve the binding affinity to SARS-CoV-2. They minimized (protonated, if needed) the Protein Data Bank (PDB) structures (Burley et al., 2017) using the “Molecular Operating Environment” (MOE) program (Vilar et al., 2008), and continued with ROSETTA (Ó Conchúir et al., 2015; Weitzner et al., 2017) and FastRelax ROSETTA (Maguire et al., 2021) to later apply machine learning. MOE was used for residue scanning as well as two machine learning models: TopNETTree (for local geometry of protein complexes) and SAAMBE3D (for a variety of chemical, physical as well as sequential, and mutation properties). Also, the z-scores were calculated using median and median absolute deviations. They used CR3022 and two other mAbs (m396 and 80R) as templates for their diversified epitopes, complexed with SARS-CoV-2 RBD. Their results suggest combining these three mAbs for higher neutralization activity. Nguyen et al. (Nguyen et al., 2021) also found electrostatic interactions explaining the higher binding affinity of CR3022 for SARS-CoV-2 than the 4A8 antibody in their all-atom and CG MD simulations (including steered MD). They used the Jarzynski equality to estimate the non-equilibrium binding free energy. They analyzed H-bonds and non-bonded contacts and used Debye-Hückel’s theory to model the electrostatic interactions for both RBD and N-terminal domain (NTD) binding sites containing charged residues. Their results indicate that effective Ab candidates should contain many charged amino acids in the regions binding to spike protein. As the for binding important residues in spike protein are positively charged (Lys and Arg), the mAbs should correspondingly contain negatively charged residues (Asp and Glu) as anticipated before (Corrêa Giron et al., 2020). Marti et al. (Martí et al., 2021) have applied classical MD simulations and accelerated MD (aMD) for enhanced sampling. They have included the complexes between the RBD of SARS-CoV-2 spike (S) glycoprotein and CR3022 or S309 antibodies and the ACE2 receptor. From MD, they calculated the potential of mean force to obtain the free energy profiles for the complexes with the RBD. They also use QM/MM for selected snapshots. With their protocol, they could explore a large part of the conformational space. They found the affinity in protein- protein complexes to follow the decreasing order: S/CR3022 > S/309 > S/ACE2. Shariatifar and Farasat (Shariatifar and Farasat, 2021) have also performed MD simulations for SARS-CoV-2 RBD complexed with CR3022 and its modifications and calculated the free binding energies. They use the FastContact software to select mutations favorable for the wild type to produce two variants of CR3022 based on their amino acid binding conformations, showing a clear affinity enhancement compared to the wild type.

### Present work framework

Some of the methods mentioned above were combined in this work (see Figure 1), specifically: (**a**) a structural-bioinformatics-based methodology to explore macromolecules as potential candidates for a higher RBD affinity using an existing experimental RBD-CR3022 complex as a template (steps 1 and 2 in Figure 1), (**b**) classical MD simulations combined with an enhanced sampling to precisely quantify the free energy of interactions for the RBD-binder complexation (steps 3 and 5 in Figure 1), and (**c**) a quicker MC sampling of a more simplified protein-protein model to investigate a larger number of complexes and search towards an optimal binder (steps 4, 7 and 8 in Figure 1). Additional atomistic simulations were also performed to explore more details of the Ab-Ag interface (step 6 in Figure 1). Such a combination of tools allows us to explore different aspects of the complex physical interactions involved in the Ab-Ag complexation process. The pros and cons of these approaches, together with the procedure in which they were combined here, are presented below. Being the Ab-Ag complexation at least a three-step process [(**i**) approximation of the two macromolecules (long-range interaction), (**ii**) structural rearrangements at short-range separation, and (**iii**) the “lock” phase at the close-contact level], (Wade et al., 1998) the combination of these tools is useful to investigate all the involved phases. All the present studied systems do not incorporate the conformation of the transition state between the open and closed states of the spike protein. It is assumed that the RBD is already available to interact with the other biomolecules (Corrêa Giron et al., 2020).

**Figure 1:**
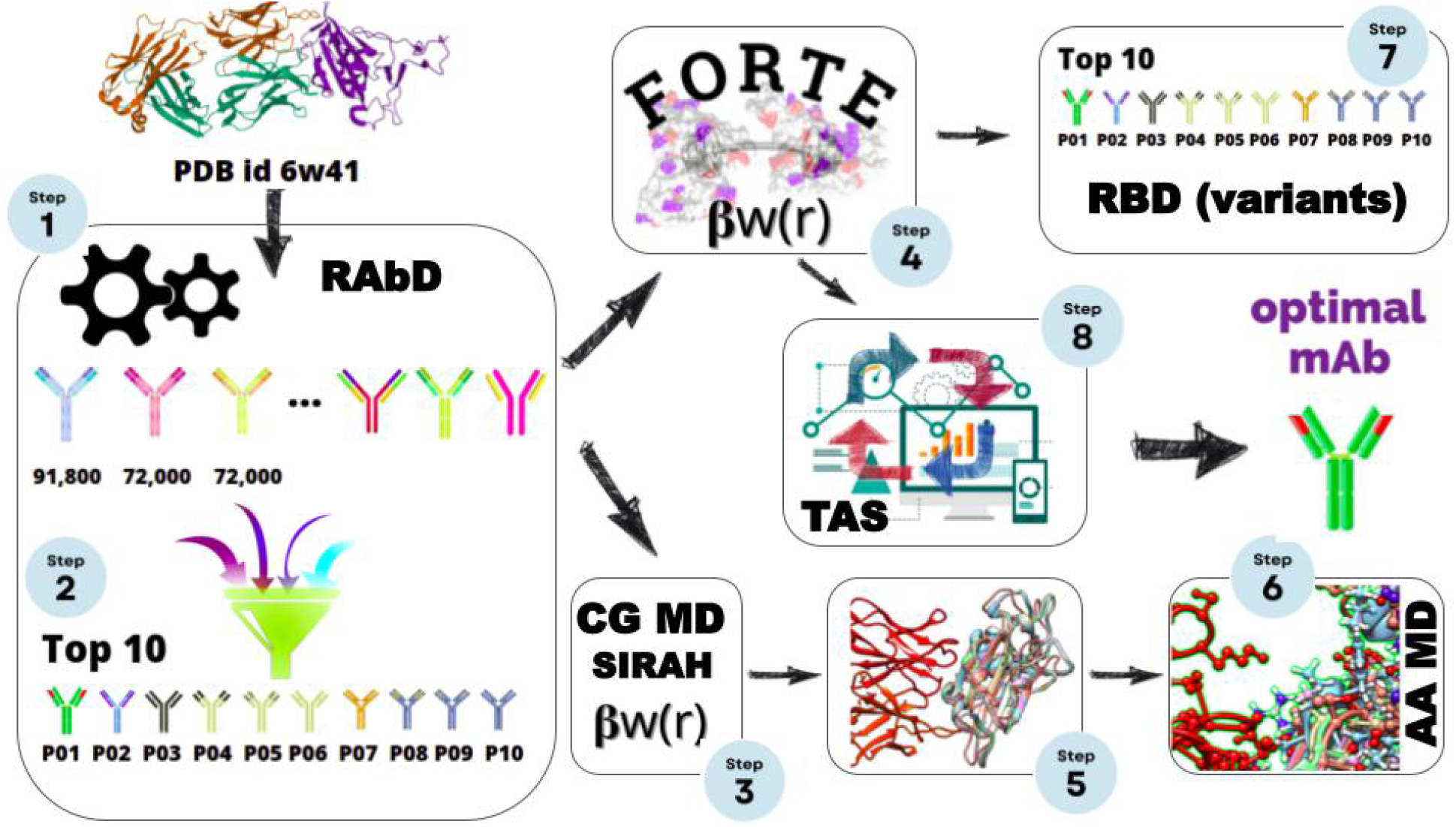
Scheme for the multiple scales *in silico* protocol, consisting of an initial structural-bioinformatics-based methodology to explore macromolecules as potential candidates (steps 1 and 2), constant-charge CG MD (steps 3, and 5), a constant-pH CG MC simulations (steps 4, 7, and 8), and an atomistic constant charge MD simulation. At the end of this cycle, an optimized mAbs with a higher binding affinity is obtained. See the text for more details.

### Structural-bioinformatics-based approach

#### RAbD approach

The RosettaAntibodyDesign (RAbD) (Adolf-Bryfogle et al., 2018) tool was employed as the first step to suggest potential binders candidates for the next phases (see Figure 1). The RAbD protocol consists of alternating outer and inner Monte Carlo design cycles. Each outer cycle consists of randomly choosing CDRs (L1, L2, L3, H1, H2, H3) from clusters in the RAbD database and then grafting that CDR’s structure onto the antibody framework in place of the existing CDRs (GraftDesign). The program then performs *N* rounds of the inner cycle, consisting of sequence design (SeqDesign) followed by energy minimization. Each inner cycle introduces mutations and structurally optimizes the backbone and repacks side chains of the CDR chosen in the outer cycle to optimize interactions of the CDR with the antigen and other CDRs. Considering that any antibody has 6 CDR’s (i.e. L1, L2, L3 on the light chain and H1, H2, H3 on the heavy chain) one has to decide which of these CDR’s should be modified with respect to the original structure (CR3022 as given by PDB id 6w41). We ran 3 sets of calculations: (**A**) all CDR’s were considered for design, subjected to SeqDesign (Adolf-Bryfogle et al., 2018); (**B**) all CDR’s except L1 were considered for design. L1 in the original CR3022 antibody is an extended loop that makes a large surface area contact with the RBD antigen (wildtype/Wuhan sequence) which stabilizes the Ab-Ag interaction. So, L1 was subjected only to sequence design, not graft design as the latter one also modifies the length of the CDR; (**C**) all CDR’s except L1 and H3 were considered for design. L1 was modeled as in step (B). H3 is located in between the H and L chains. From all the 6 CDR’s, H3 is not canonical, i.e. it does not adopt classifiable conformations (clusters of conformations). When visually analyzing the RAbD generated conformations, it became obvious that some candidates with both very high scores and Ab-Ag interface surface areas were not realistic **(Figure S1)**. For example, large Ab-Ag interface surface areas encountered in some cases were due to very long H3 CDRs. However, care should be taken when predicting H3 CDR, especially for long ones. Thus, H3 was not subjected to graft or sequence design in this scenario. Poses from all three sets of calculations were considered for further evaluations, manually excluding the unrealistic ones. In total, we generated a number of 91,800 candidates for (**A**), 72,000 candidates for (**B**), and 72,000 candidates for (**C**) scenarios.

From the entire pool of candidates, we have generated a comprehensive list of potentially improved affinity Ab-Ag complexes using two criteria: (**I**) the complex must have an Ab-Ag interface score below -150 REU (ROSETTA Energy Units) (the native interface score is -65 REU), and (**II**) an Ab-Ag interface surface area larger than 1900 Å^2^ (the native complex interface surface area is 2060 Å^2^). The interface surface area is important as a larger total surface area means a higher specificity due to shape complementarity. Similarly, a larger hydrophobic contact area means a higher affinity while a larger polar contact area means a higher specificity. There were 41 Ab-Ag complexes that fulfilled the imposed criteria **(Table S1)**. According to the ROSETTA scoring function, all these antibody-antigen complexes present better affinities for SARS-CoV-2 RBD (wild type sequence) than the original CR3022 antibody. The first 10 best candidates (P01 to P10) from the final RAbD list plus the native complex were considered for the next analyses.

### Coarse-grained molecular dynamics (MD) with enhanced sampling approach for free energy calculations

Umbrella sampling (US) constant-charge MD simulations were employed to evaluate the free energy of binding of the top ten Ab-Ag complexes selected from the above described RAbD calculations (Step 3 in Figure 1). For the efficiency of calculations, a reduced representation of the interacting partners was adopted, using the SIRAH 2.2 coarse grain (CG) force field (ff) (Machado et al., 2019). This approach was successfully used in the past for estimations of free energy of binding in the case of Ab-Ag complexes (Patel and Ytreberg, 2018). During the US procedure, the geometry of the mAb was restrained by applying weak harmonic position restraints (20 kJ mol^−1^nm^−2^ force constant) on the CG beads corresponding to the protein backbone. The distance between the center of mass (COM) of mAb and the RBD was considered as the reaction coordinate. Cylindrical positional restraints (as defined in the GROMACS 2019 suite (Abraham et al., 2019, 2015)) were applied on the RBD to allow for its movement relative to mAb along the reaction coordinate. A total number of 36 windows were used for US simulations which ensured a sufficient histogram overlapping for accurate results. Within each window, a US potential with a force constant of 1500 kJ mol^−1^nm^−2^ was applied to the COM of the RBD. Ab-Ag complexes were simulated for 16ns in each US window, the first 2ns being excluded from the analysis. For each Ab-Ag candidate, US simulations were repeated 10 times, which allowed for results averaging and errors estimation. The Potential of Mean Force (PMF) profiles were constructed using Weighted Histogram Analysis Method (WHAM) (Hub et al., 2010). All simulations have been performed with the Wuhan sequence (wild-type) at constant temperature (300K) and pressure (1 atm) using GROMACS 2019.3 suite (Abraham et al., 2019, 2015) on Beskow supercomputer at PDC Stockholm, Sweden. The initial structure was retrieved from the Research Collaboratory for Structural Bioinformatics PDB with the id 6w41 (X-ray data with a resolution of 3.08Å, pH 4.6) (Yuan et al., 2020). This set of simulations was carried out at a constant-charge condition assuming pH 7. Internal degrees of freedom were present in the model to allow for the molecular conformational adjustments upon the binding process.

### A fast constant-pH coarse-grained (CG) approach for free energy calculations on a large scale

The same ten selected fragments of mAbs candidates obtained from the RAbD analysis were also submitted to exhaustive investigations utilizing a fast constant-pH CG biophysical model specially designed for protein-protein complexation (Barroso da Silva et al., 2016; Delboni and Barroso da Silva, 2016; Kurut et al., 2012; Persson et al., 2010) – see step 4 in Figure 1. These cost-effective CG simulations are less expensive than the US calculations with the CG SIRAH ff described above and have the benefit to allow the description of amino-acid charge variations (due to protonation state variations) occurring at constant pH. This implies that important electrostatic interactions as the charge regulation mechanism are properly included in the model (Adžić and Podgornik, 2015; Barroso da Silva et al., 2018, 2006; Barroso Da Silva and Jönsson, 2009; Lund and Jönsson, 2013). This approach is aimed to capture only the main features of the complexation phenomena with a clear emphasis on the electrostatic interactions. Different studies have highlighted the importance of these interactions for the host-pathogen and antigen-specific antibody interfaces (Bai and Warshel, 2020; Corrêa Giron et al., 2020; Nguyen et al., 2021, 2020; Poveda-Cuevas et al., 2020, 2018; Xie et al., 2021). Successful applications of this simplified model to several biomolecular systems have been previously reported in the literature including viruses proteins and the SARS-CoV-2 RBD-mAb interactions (Barroso da Silva et al., 2016; Corrêa Giron et al., 2020; Delboni and Barroso da Silva, 2016; Kurut et al., 2012; Lund et al., 2008; Mendonça et al., 2019; Persson et al., 2010). We shall refer to this electrostatic model as FORTE (**F**ast c**O**arse-grainedp**R**otein-pro**T**einmod**E**l). The core of such a model is the fast proton titration scheme (FPTS) (Barroso da Silva and MacKernan, 2017; Teixeira et al., 2010) combined with the possibility to translate and rotate the macromolecules using the Metropolis MC method (Metropolis et al., 1953). Charged and neutral spherical beads of different radii mimicking titratable and non-titratable amino acids, respectively, interact via Coulombic and van der Waals terms (Barroso da Silva et al., 2016; Delboni and Barroso da Silva, 2016). Protein coordinates given by the structural-bioinformatics-based approach were directly converted into this amino acid model. Heteroatoms and the solvent were removed from the input structures.

Early ideas of such a model are rooted in the works of Marcus (Marcus, 1955) and Jönsson and co-authors (Barroso da Silva and Dias, 2017; Jönsson and Svensson, 1993; Svensson et al., 1990). The reduction in the degrees of freedom together with the description of the proteins at the mesoscopic level in a continuum solvent model is a clear advantage from a computational point of view. For instance, the smaller amino acid (glycine) is modeled by four sites in the SIRAH ff instead of a single one as adopted in FORTE. The reduction is even more significant for larger residues as aspartic acid where the drop is from 11 to 1. This can result in a decrease of the computing time by a factor of 11^2^ making it possible to apply it on a large-scale scenario. On the other hand, the main drawbacks are the assumption of a rigid body description to model the macromolecules and the ambiguity involved in the choices of the van der Waals contributions (Barroso da Silva et al., 2016; Corrêa Giron et al., 2020). Yet, to form the RBD-Ab complex, the two molecules have to come closer before conformational changes happening at short-range separation distances can be important to add additional attraction and/or stabilize the formed complex (Wade et al., 1998). Furthermore, these possible limitations have proven to be not critical in several cases studied before. The present outcomes also contribute in this direction as discussed in the results section below. More details of this electrostatic protein-protein model are given elsewhere (Barroso da Silva et al., 2016; Corrêa Giron et al., 2020; Delboni and Barroso da Silva, 2016). All calculations with FORTE were performed here at pH 7, 150mM of NaCl, and 298K. After equilibration, at least 3×10^9^ MC steps were run during the production phases. Three replicate runs were employed for each simulated system. They were also used to estimate the corresponding standard deviations.

The main quantity of interest extracted from these biophysical simulations was the free energies of interactions, or PMF [*βw(r)*, where *β*=1/K_B_T, K_B_ = 1,380×10^−23^ m^2^ kg s^−2^K^−1^ is the Boltzmann constant, and T is the temperature, in Kelvin], as a function of the macromolecules separation distances [*r*]. They were directly calculated from their center-center pair radial distribution functions [*βw(r)=-ln g(r)*] and sampled using histograms during the production phase of the MC runs. For long enough simulations, *βw(r)* is typically obtained with good accuracy and is able to reproduce experimental trends (Corrêa Giron et al., 2020). The relatively lower computation costs in comparison with other theoretical approaches allow the repetition of the calculations at different physical-chemical conditions (e.g. different solution pHs) and macromolecular systems (e.g. RBDs with several different mutations) using high-performance computers. This is a key aspect to investigate the binding affinities for a set of several RBDs (with all possible mutations of interest), different binder candidates and to perform further optimization of the best ones. A large number of runs are needed which are prohibitive with more elaborated molecular models.

Different sets of calculations were carried out with FORTE. Initially, the binding properties of the 10 best binder candidates (protein P01 to P10) that were obtained from the structural-bioinformatics-based analysis (using RAbD) were tested to form complexes with the RBD from the wild type (wt) SARS-CoV-2 (Wuhan sequence): RBD_wt_+Px → RBD_wt_Px, where *x* ranges from 1 to 10 (i.e., the ten top candidates). *βw(r)* was calculated for all systems. Simulations with the fragment of the original mAbs CR3022 (P00) were also performed for comparison. All these simulations were repeated for RBDs built up with sequences with different mutations of interest present in reported variants (see step 7 in Figure 1): (a) N501Y (Alpha/B.1.1.7), (b) K417N, E484K and N501Y (Beta/B.1.351), (c) L452R, T478K and E484Q (Delta/B.1.617.2), (d) L452R (Epsilon/B.1.427/B.1.429), (e) E484K (Eta/B.1.525), (f) K417T, E484K and N501Y (Gamma/P.1), (g) E484K and N501Y (Iota/B.1.526 NY), (h) L452R and E484Q (Kappa/B.1.617.1), (i) G339D, S371L, S373P, S375F, K417N, N440K, G446S, S477N, T478K, E484A, Q493R, G496S, Q498R, N501Y, Y505H (Omicron/B.1.1.529), and (j) Y453F (mink) (Annavajhala et al., 2021, p. 526; Callaway, 2021; Deng et al., 2021; Ford et al., 2021; Khateeb et al., 2021). The input structures with the mutations in the SARS-CoV-2 RBDs were prepared with “UCSF Chimera 1.15” (Pettersen et al., 2004) by the simple replacement of the amino acid followed by a minimization with default parameters and considering the H-bonds (Pettersen et al., 2004). Structures obtained by this simple procedure seem to be equivalent to other available ones (Corrêa Giron et al., 2020). For instance, a comparison between a recent RBD structure predicted by AlphaFold2 for the Omicron variant (Ford et al., 2021) with the one generated using “UCSF Chimera 1.15” gives a root-mean-square difference of 0.5Å. This suggests that the mutations seen so far do not have a significant effect on the overall folded structure of the RBD.

After confirming the outcome from the US calculations with FORTE, an optimization procedure was invoked to explore the possibility to further improve the binding features of the fragment of mAbs given by RAbD (step 8 in Figure 1). The two best binders (P_best_ and P_best-1_) with a higher chance to block the RBD_wt_were further submitted to a “theoretical alanine scanning” (TAS) (Corrêa Giron et al., 2020), a technique used here to determine the contribution of a specific amino acid to the RBD-binding. In this process, an amino acid from a given binder was replaced by ALA, and the complexation simulation was repeated for this new possible binder. This mutation was done directly at the mesoscopic level of the proteins without the need to pre-generate its coordinates by the above-described procedures. It is assumed that the tested point mutations will have a minor effect on the overall folded structure of the mAb. The minima values of *βw(r)* for each new system (RBD_wt_ interacting with P_best_ or P_best-1_ carrying the ALA mutation) are recorded for comparison after all possible single replacements were fully explored one by one. This theoretical lead optimization protocol is schematically illustrated in Figure 1.

A three-cycle process was followed in this optimization pipeline for mAb engineering with higher affinity. After all possible ALA substitutions were tested one per time, the best single mutation obtained from this first cycle was incorporated into the binder resulting in a binder’. This new protein binder’ contains a single ALA mutation in a specific position that resulted in the higher RBD_wt_ affinity among all tested possibilities. The process was restarted with this new macromolecule (i.e. binder’) being subjected to another TAS loop. After this second loop, the binder’ has a new amino acid substitution in its sequence together with the previous one already incorporated by the ALA single mutation in the first cycle (i.e. at this point, the binder’’ has two ALA replacements [A-A]). A third cycle of TAS was also done using the binder’’ with the double ALA mutation to explore the effect of an additional ALA replacement in its sequence (A-A-A). Besides TAS, equivalent tests were also done with the substitution of any residue by GLU in a “theoretical glutamic acid scanning”. This acid residue was chosen based on a previous work where we could observe some dependency on an increase in the binding affinities with a decrease in the total net charges of the binders (Corrêa Giron et al., 2020). This process with three cycles was repeated for GLU as done with ALA. All these replacements by either ALA or GLU were combined in different ways (A-A-A, A-A-E, A-E-E, and E-E-E). Each possible combination was evaluated to guide the selection of the systems that improved the binding affinity for the RBD_wt_. Only up to three simultaneous substitutions in the wildtype mAb were tested to avoid a complete mischaracterization of the original template protein (P_best_ or P_best-1_). This could result in a complete unfolding of the macromolecule and/or an increase in the chances for Ag-Ab aggregation. Moreover, at this point, the number of different simulated systems was too large, and the computational costs started to be prohibitive even for a relatively cheaper CG model.

Finally, the designed protein from these three engineering cycles (P’_best_) with the higher RBD_wt_binding affinity was tested with the other RBDs from the main variants of concern and interest. Additionally, titration simulations employing the FPTS were used to provide the main physical-chemical properties of the individual macromolecules (all from the wild-type P00 to P10 and each new macromolecule produced by the binding optimization technique with a single, double and triple mutation). In these runs, the total net charge numbers of these binders were computed in the absence of the RBD, i.e. the FTPS was employed for a single binder in an electrolyte solution. Such a large set of net charges data was used to further investigate the possible correlations between the binding affinities and the mAb’s net charge. This was useful to complement the previous initial analysis performed with a quite smaller number of pairs of RBD-binders in comparison with this larger data set (Corrêa Giron et al., 2020).

### Atomistic MD simulations

Conventional MD simulations of the wild-type CR3022-RBD (PDB id: 6w41) and P01-RBD (RAbD generated) complexes have been performed using the Amber99SB-ILDN ff (Lindorff-Larsen et al., 2010) for protein description and the TIP3P (Jorgensen et al., 1983) model for water (step 6 in Figure 1). The Ab-RBD models have been solvated with a sufficient number of solvent molecules to ensure a minimum distance between periodic images of 2.2 nm. Ions have been added to neutralize the solvated systems. The length of the simulations was 50ns at constant pressure (1 bar) and temperature (298K). The GROMACS 2019.3 suite (Abraham et al., 2015) has been used for all atomistic MD simulations.

## RESULTS AND DISCUSSION

### ROSETTA-designed antibody structure: identifying the top best binder candidates

The general workflow of the antibody design procedure consisted of generating a large number of Ab-Ag structures using RosettaAntibodyDesign software (Adolf-Bryfogle et al., 2018), followed by ranking of the generated models according to the ROSETTA scoring function. In short, the RAbD algorithm is meant to sample the diverse sequence, structure, and binding space of an antibody-antigen complex. We have generated 235,000 candidates who were ranked according to the ROSETTA scoring function. From the total number of designed candidates, only 383 showed better affinities compared with the native CR3022-RBD complex (Figure 3).From this list, we choose 10 complexes (according to the criteria described in the “Theoretical Methods” section) that were further evaluated by free energy (ΔG) calculations using both umbrella sampling (US) and FORTE (CDRs primary sequences are presented in Figure 4). We then considered, as the outcome of the computational study, the structures that give consensus results in all three types of evaluations. Considering that any computational method has its limitations and accuracy, evaluating the Ab-Ag complexes with complementary computational approaches, and selecting the ones that are consistent in all evaluations, strongly increase the reliability of predictions. Thus, the observed improved affinity of the models could be considered to result from the real physics/chemistry of the complex interactions in the systems and not from any particular algorithm-related issues.

**Figure 2:**
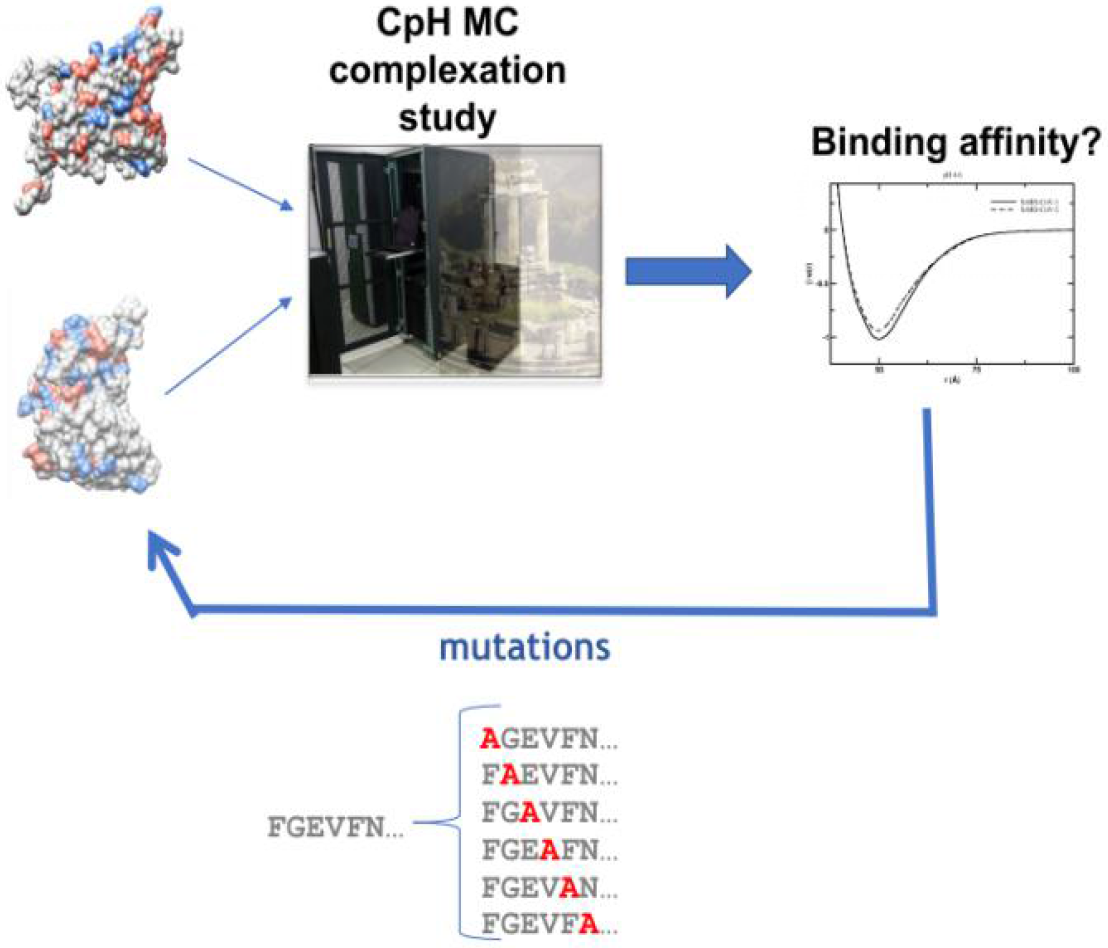
Scheme for the procedure used for the theoretical electrostatic mAb optimization using TAS. A pair RBD-mAb is submitted to a constant-pH complexation study using FORTE where the two macromolecules can titrate, translate and rotate in all directions. After each simulation run, the free energy of interaction is saved for comparison at the end of the full cycle. A substitution of an amino acid by ALA is introduced in the wild-type mAb. A new complexation run is carried out with this new binder (i.e. the wild type with a new single ALA mutation). The process is systematically repeated for all residues. Only one residue is replaced by ALA per time. This procedure corresponds to step 8 in Figure 1.

**Figure 3.**
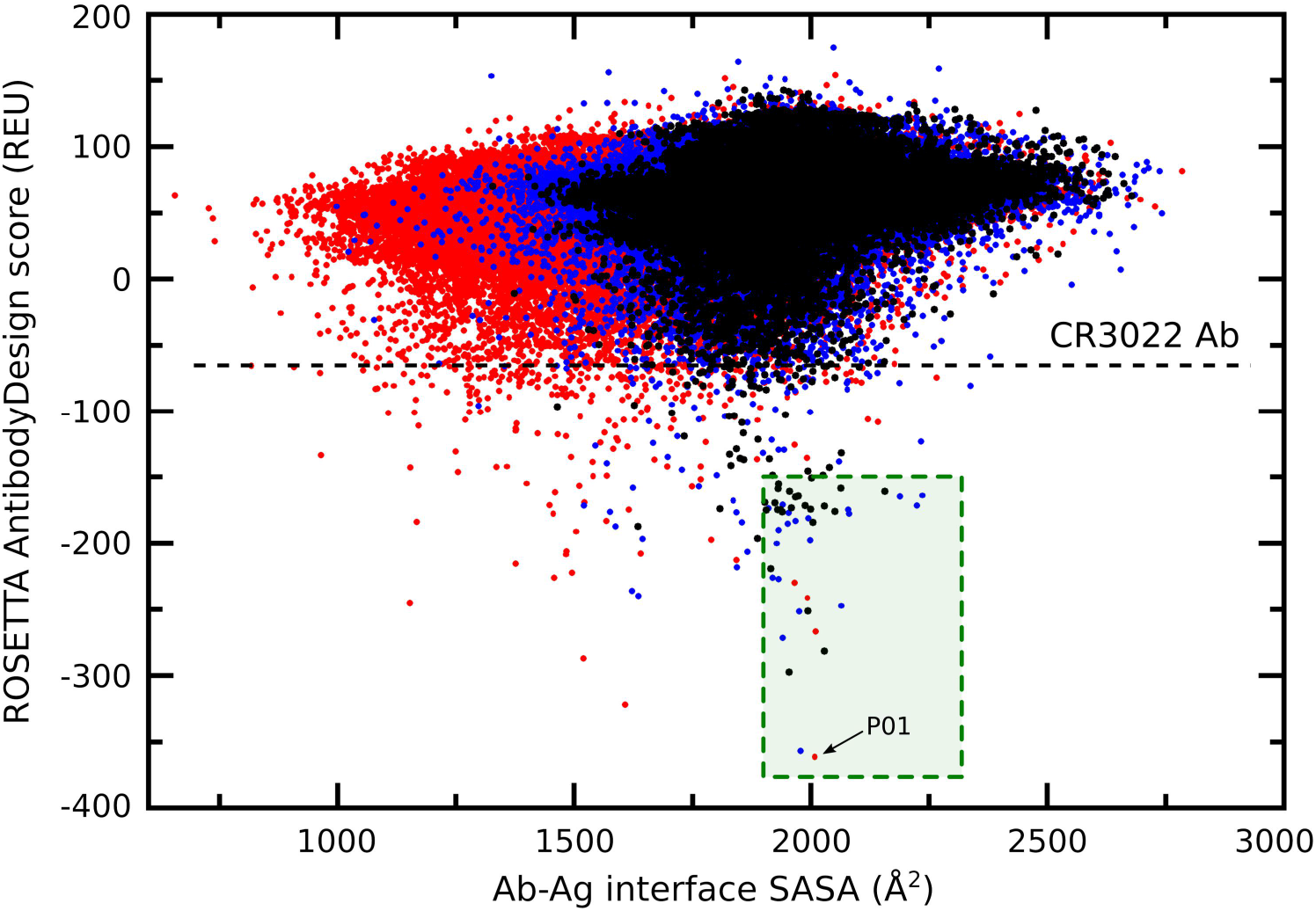
2D plot of RAbD score against antibody-antigen interface SASA (solvent accessible surface area). The horizontal line corresponds to the CR3022/RBD complex RAbD score given in Rosetta Energy Units (REU). The green square indicates the area in the two dimensional plot that corresponds to the mAb candidates selection criteria.

**Figure 4.**
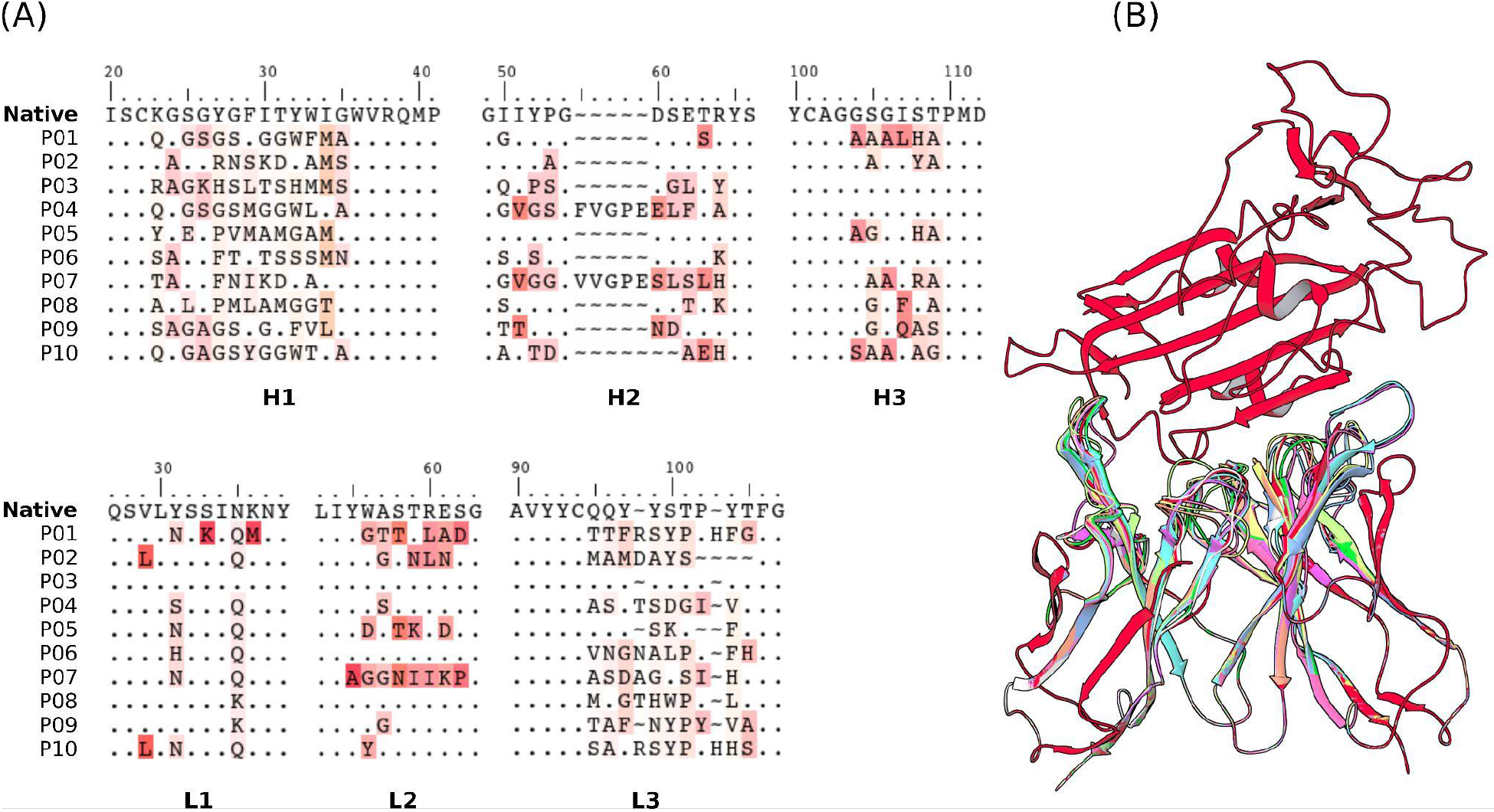
Sequences of the CDRs (A) and the structures (B) of the 10 selected mAbs for evaluations using US/SIRAH and FORTE methods. Dots represent identities. Residue letters in (A) were colored according to similarity with their counterparts in the reference native CR3022 sequence. In (B) RBD (up) was colored in red and each candidate antibody (down) in a different color.

### RBD-Ab free energy of interactions explored with umbrella sampling CG simulations

Free energy calculations using US MD calculations revealed that from the list of the top 10 best RAbD candidates only P01, P05, P06, P09, and P10 showed better binding affinities compared with the native CR3022 (see Figure 5 for their PMF profiles). Although all these candidates show statistically significant improved affinities compared with CR3022, the errors in estimating the binding free energies for P05, P06, P09, and P10 did not allow us to distinguish between them in terms of affinity. Among the candidates P01 seems the best, giving statistically significant better affinity of binding compared with both the native CR3022 and the P05, P06, P09, and P10 set.

**Figure 5.**
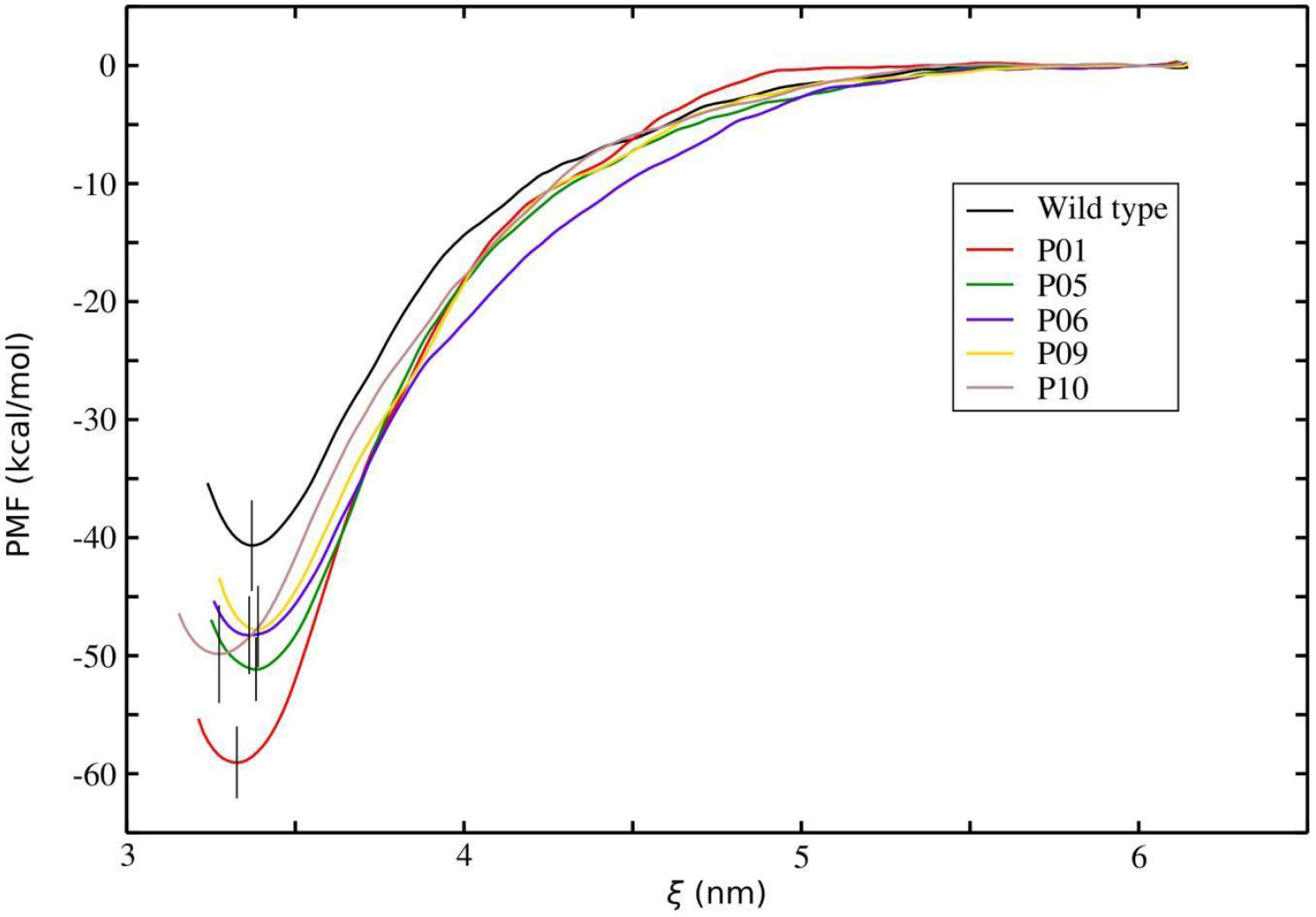
Averaged RBD-Ab PMF profiles for the wild type and the candidates that give statistically significant better binding affinity in the constant-charge MD simulations with the SIRAH CG force field using the umbrella sampling method. The vertical bars represent the estimated standard deviations of the minimum of the PMF profiles. See the text for more details.

However, caution must be taken when considering absolute values obtained by US-CG SIRAH to compare with more rigorous atomistic or experimental data. As the results of Patel et al. (2017) showed, only the ΔΔG = ΔG_mutant_ -ΔG_wild-type_ should be regarded quantitatively, as the granularity of the SIRAH-CG model may affect the absolute values of the calculated ΔG from the PMF profiles. On the other hand, ΔΔG computations based on SIRAH-CG did fit well with experimental data (Patel and Ytreberg, 2018) and are much more efficient than atomistic US approaches, which of course give more precise absolute values for ΔG (Nguyen et al., 2021). Therefore, it is meaningless to use the CG-US results for a direct comparison with experimental data like K_d_.

The free energy evaluations based on CG-US showed ΔΔG values of -18.3 kcal/mol for P01 and -7.1 kcal/mol to -10.8 kcal/mol for P01-P05-P06-P09-P10 group relative to the wild-type CR3022 antibody. These values are comparable to experimental kinetic data found in the literature for stabilizing mutations induced in Ab-Ag complexes (Moal and Fernández-Recio, 2012; Pons et al., 1999; Pruett and Air, 1998).

### RBD-Ab free energy of interactions explored with FORTE

Despite its approximations, the simplified CpH CG protein-protein model is able reproduce the main trends obtained from using the more elaborate models (see above). Figure 6 shows the minimum values of the free energy of interactions [*βw_min_*] for the complexation between the SARS-CoV-2 RBD_wt_ with the fragment of CR3022 and with all engineered proteins given by RAbD from P01 to P10. These new binder candidates all have a higher RBD_wt_affinity as observed for the simulations with SIRAH ff too (see Figure 5). They can be ranked as P01/P06 > P02/P05/P08/P09/P10 > P04 > P03 > P05 in terms of their binding affinities. It is virtually impossible to distinguish the proteins within the groups P01/P06 and P02/P05/P08/P09/P10 due to their estimated standard deviations. Conversely, there is no difficulty in determining that the two best candidates are P01 and P06. Both proteins have virtually identical free energy profiles for their complexation with the RBD_wt_ [-0.788(6)K_B_T and -0.790(6)K_B_T for P01 and P06, respectively). Yet, P01 was selected as P_best_ to follow the consensus with the best protein found in the previous biophysical simulations with the SIRAH ff (see Figure 5). Since it is not exactly the same physics that is incorporated into the two models, this consensus policy reinforces that P01 has special features for a stronger binding with the RBD_wt_.

**Figure 6:**
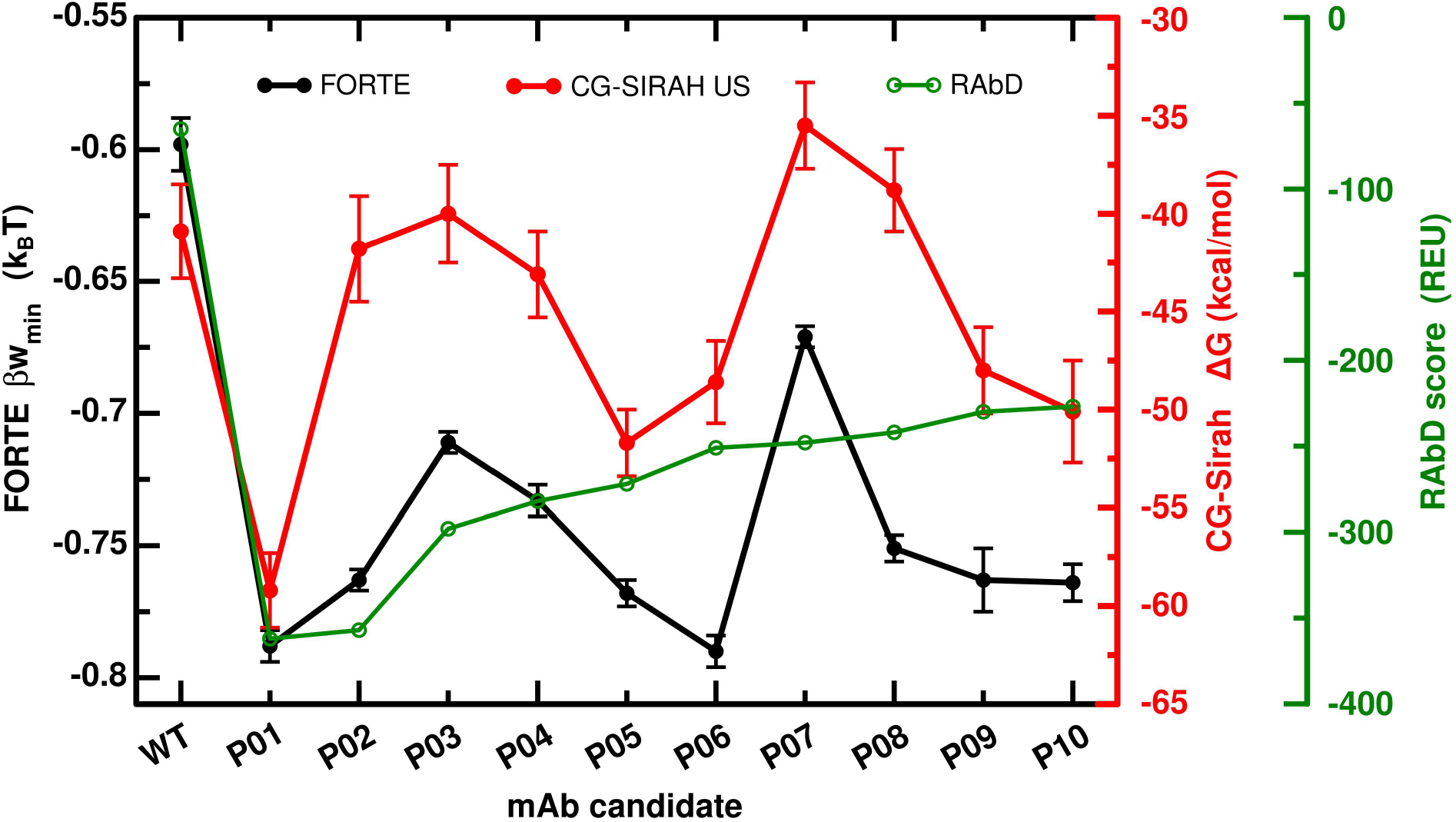
Minima free energy of interactions values [*βw_min_*] measured for the SARS-CoV-2 RBD-mAbs complexation at pH 7 and 150mM of NaCl by the CpH MC simulations (FORTE) for the wildtype RBD_wt_ (Wuhan sequence) and different fragments of mAbs candidates (P01 to P10). The reference (P0) is the data for the complexation with the native fragment of CR3022 (from the PDB id 6w41). The error bars were calculated using the three replicates for each simulation system. The minima values obtained by the classical MD with SIRAH ff as given in Figure 5 are included in the figure for the sake of comparison. Values from the RAbD in REU are also reported for comparison.

Although P06 was not found equivalent to P01 in Figure 5, it could still be an interesting candidate. Therefore, P06 was treated as P_best-1_. The fact that FORTE provided these two binders (P01 and P06) with equivalent binding affinities instead of a single one as seen in the previous approach could be due to a couple of reasons: **a**) a smother energetic landscape in this CG model eventually not able to describe the same roughness of the phase space, **b**) the presence of the charge regulation mechanism in the FORTE approach (absent in the constant-change simulations) that give rise to an extra attraction, **c**) the lack of structural rearrangements in the FORTE approach. Nevertheless, even in the US/SIRAH ff approach, the minima values suggest a binder better than the native CR3022 (see Figure 5).

An interesting comment here is the fact that groups of binders do share equivalent RBD binding features. This suggests that mAbs produced from previous infections or by the seroconversion of a given vaccine could have a chance to neutralize more than one strain in agreement with clinical reports (Edara et al., 2021). It also illustrates that a weaker response can be obtained depending on the specific mutations as also expected. Equivalent reasoning can be done for therapeutic molecules designed to prevent the attachment of the spike protein via the RBD in the human cell. For instance, Wang and co-authors described a potent mAb from covalent patients that works well against 23 variants (Wang et al., 2021).

It is worth reminding that a limitation of this CG model is the precise quantification of the absolute numbers in the reported values for the PMF. It is especially difficult to quantitatively (not qualitatively) reproduce experimental K_d_ values and/or typically *βw(r)* values obtained by more detailed force field descriptions (Barroso da Silva et al., 2016; Corrêa Giron et al., 2020; Delboni and Barroso da Silva, 2016). The same problem described above for SIRAH ff might be more severe for FORTE due to its higher granularity. Yet, any ambiguity in the interpretation of the obtained data can be solved when results are interpreted in relative terms. For instance, comparing the measurements between a set of similar macromolecules under the same experimental conditions (Barroso da Silva et al., 2016; Corrêa Giron et al., 2020). This is exactly the case for this comparison between different pairs of RBDs and a putative binder. Both CG approaches are useful for distinguishing between stabilizing and destabilizing CDR modifications, for ranking different candidates relative to the wild-type and in-between them. Qualitative comparison can still be made as shown in Figure 6.

After validating the theoretical predictions obtained with FORTE by means of the reproductivity of results from the other methods, we expanded its analysis and tested the binding affinities of these Rosetta-designed fragments of mAbs to other SARS-CoV-2 RBDs with critical mutations. It is particularly interesting to assess if the engineered proteins are good binders also for the present VOCs (Alpha, Beta, Gamma, Delta, and Omicron) and the variant of interests (VOIs). Such information would be vital for any practical therapeutic applications of these mAbs in the future. The free energy data at the same experimental conditions as in Figure 6 for several pairs of RBDs and mAbs was compiled and displayed as a heatmap-style plot in Figure 7. In this Figure, the darkest blue is the case with the highest binding affinity. Conversely, the darkest red is given to the system with the lowest binding affinity. This is observed for the native CR3022 (P00) interacting with any studied RBD. All putative Rosetta-designed binders improved the binding features in comparison with the native CR3022. As it can also be seen, the general behavior for all RBDs is similar to what was observed for the interaction with the SARS-CoV-2 RBD_wt_(see Figure 6). All molecules P01 to P10 have the potential to block the interaction between virus and host cell preventing the RBD to be available to anchor ACE2. The two best candidates for all these RBDs identified by FORTE were also P01 and P06. This reinforces the conclusions concerning their potential ability to neutralize both the wild-type SARS-CoV-2 virus and the main present variants. Considering the estimated error bars, all Rosetta-designed proteins follow the same trend seen in Figure 6 for RBD_wt_. They also tend to respond with similar affinities to all studied RBDs, i.e., a good binder for one specific RBD is a good binder for another RBD too. However, there are interesting exceptions. For example, P03 has a relatively lower affinity for RBD_Delta_, RBD_Eta_, RBD_Kappa_, and RBD_mink_ in comparison with the RBD_wt_. Surprisingly, both the native CR3022 and all Rosetta-designed binders show a higher affinity for RBD_Omicron_. This suggests that both CR3022 and its derived-mAbs have a good chance to neutralize the Omicron variant.

**Figure 7:**
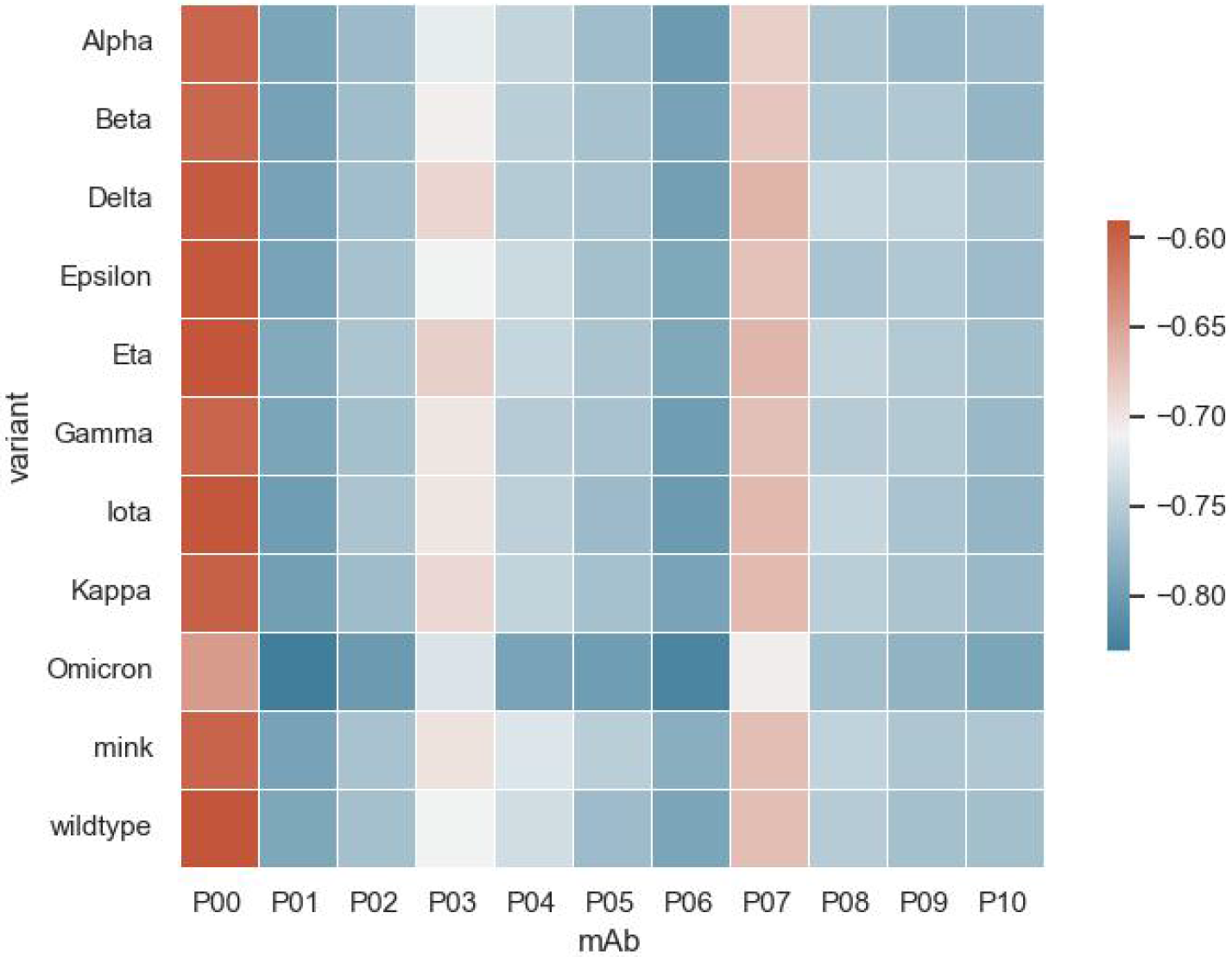
Heatmap with the minima free energy of interactions values measured for the SARS-CoV-2 RBD-mAbs complexation at pH 7 and 150mM of NaCl by the CpH MC simulations (FORTE) for the RBD of the main critical variants and different Rosetta-designer binder candidates (P01 to P10). The reference (P00) is the native fragment of CR3022. All values of *βw_min_* are given in K_B_T units. The maximum estimate error is 0.01. See the text for a description of mutations considered in each case.

### RBD-Ab local interface interactions explored with atomistic MD simulations

We will discuss in the following the structural determinants at close-contact distances, identified by atomistic simulations, responsible for the observed better affinity of P01 against RBD_wt_. Mutations designed in the P01 candidate were located in all six CDRs on both the light (L) and heavy (H) chains. These mutations improved specificity and affinity by favoring the formation of supplementary salt bridges and hydrogen bonds as well as hydrophobic interactions. Protein-protein interactions are complex processes that, at short distances, are dependent on the interface properties consisting of shape complementarity, size, short-range electrostatic interactions, polar and hydrophobic interactions. Specificity in protein-protein binding is mainly given by electrostatic interactions (Kumar and Nussinov, 2002) while non-specific and van der Waals interactions are the driving force for increased affinity (Wodak and Janin, 2002). We have used MD simulations to comparatively evaluate P01 with the wild type CR3022 in terms of different interaction contributions to RBD-Ab binding. The simulations were analyzed for the probability distribution functions (PDF) of three interface interaction descriptors (see Figure 8),namely the number of *i)* Ab-Ag salt bridges, *ii)* non-polar contacts and *iii)* hydrogen bonds (between polar-polar and polar-charged residues). Thus, for each saved MD trajectory frame the number of salt-bridges formed between charged residues of the Ab and RBD was calculated. A salt bridge was considered to exist between acidic/basic residues if the distance between the heavy atoms of the charged moieties (i.e. side-chain carboxyl and amino) was shorter than 3.5 Å. Using the entire pool (full trajectory) of calculated salt-bridge numbers a histogram was extracted with respect to the number of salt bridges (i.e. the distribution of a number of frames with a particular number of salt bridges). The histogram was then normalized considering the area under the curve to be 1 in order to obtain the PDF of Ab-RDB salt-bridges number. The number of non-polar contacts was obtained by counting, for each frame, the hydrophobic residue atom pairs (Ab-to-RBD) within a distance shorter than 3.5 Å. A similar procedure as above was applied to calculate the histogram and PDF of non-polar contacts. For polar interaction histogram and PDF, we counted the number of hydrogen bonds between polar residues and polar to charged residues in the Ab-RBD pair. It can be seen from the plots in Figure 8 that the main contributions to the increased affinity of P01 to RBD, compared with the wild-type CR3022, come from an increased number of salt-bridges and a higher hydrophobic contact area to RBD. The average number of salt-bridges at the P01 interface to RBD is ∼4 while for the wild-type CR3022 is ∼3. The Lys35 residue, which replaced a Ser35 in the CR3022, is an important player in short-range electrostatic interactions at the P01-RBD interface as it is strategically placed on the tip of the L1 loop and has the optimum side-chain length to form a strong salt-bridge with the acidic Glu in position 516 on the RBD. The average number of hydrophobic contacts between P01 and RBD (∼95) is significantly bigger than for the CR3022-RBD pair (∼125), which will result in a higher binding affinity. Four mutations in P01, that replace original residues in CR3022 with more hydrophobic ones, contribute to this increased hydrophobic contact area: W40F, S110A, G111A, I112L. They form a hydrophobic patch in the middle of the mAb – RBD interface that interacts with Lys378(4 methylene groups)-Tyr380-Pro-384-Tyr369-Val382 on the RBD counterpart.

**Figure 8.**
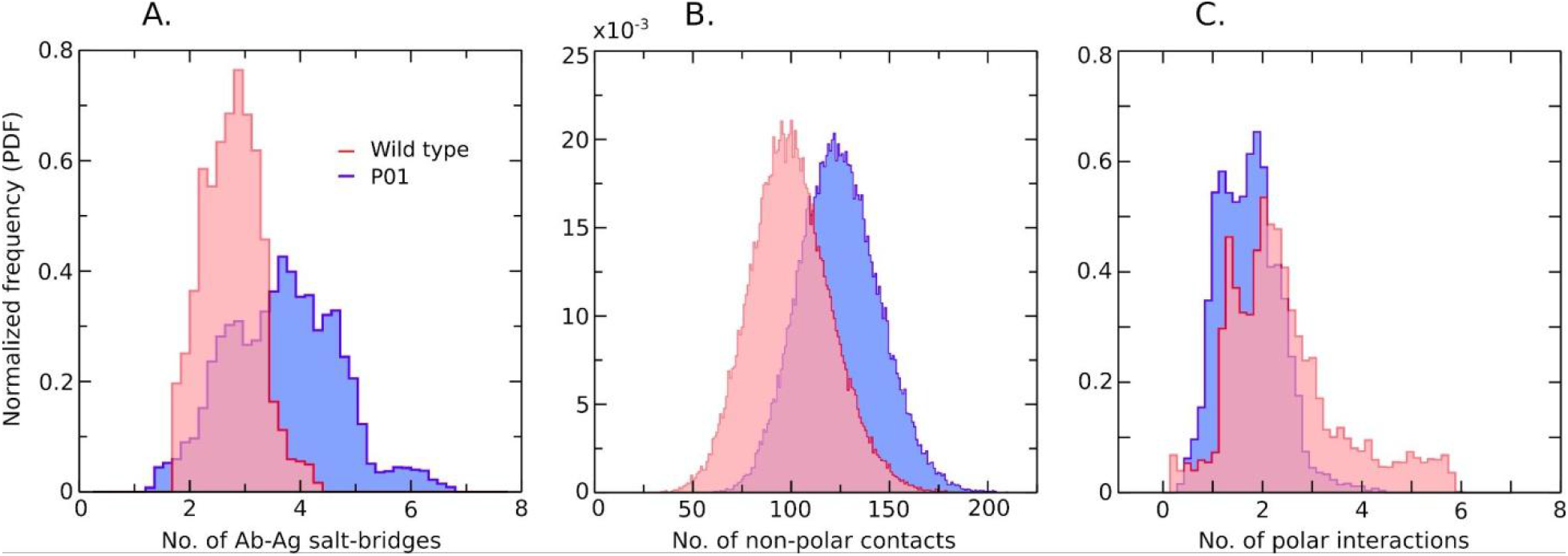
Molecular descriptors were used to analyze the contributions of different interaction types to RBD-P01 binding: normalized probability distribution functions (PFD) of Ab-Ag salt-bridges (A), non-polar contacts (B), and hydrogen bond (C) numbers.

To get a detailed picture of the RBD-P01 interaction interface, we have analyzed the impact of the introduced mutations on the local molecular environment of the complex, which is discussed below for each CDR separately.

#### L1 CDR design

The tyrosine residue in position 31 in L1 CDR was replaced by Asn in P01, which easily participates in hydrogen bonding with both Thr430 and Asp428 on RBD_wt_. The interaction of the native Tyr31 with Thr430 is impeded due to the longer side chain of tyrosine compared with asparagine. One key mutation present in L1 is S35K which replaces a neutral serine residue with a positively charged lysine. The original Ser35 residue is located on the tip of the L1 loop that penetrates deep into the Ab-Ag interface stabilizing the complex. Replacing the serine with lysine in this position provides a strong salt bridge formation between the long basic side chain of Lys35 and the acidic Glu in position 516 on the RBD_wt_, an interaction not present in the native CR3022. Replacement of Asn with Gln in position 37 provided further stabilization as the longer side chain of glutamine compared with asparagine allowed for a new interaction of Gln37 on the antibody with His519 on the RBD_wt_.

#### L2 CDR design

The replacement of the neutral hydrophilic Ser72 with an acidic Glu residue in P01 introduces a strong salt bridge formation between Glu72 and the basic Lys386 on the RBD_wt_.

#### L3 CDR design

Another mutation that increases binding specificity and affinity was introduced in the L3 loop replacing Tyr110 with Arg in P01. The substitution of neutral tyrosine with the positively charged arginine allows for a salt bridge formation with the neighboring Asp428 on RBD_wt_. The P01 designed L3 CDR has two supplementary residues Pro135 and His136 compared with the native CR3022. This makes the L3 loop more extended towards RBD_wt_, increasing the Ab-Ag contact surface area. The L3 loop also contains the mutation of Thr112 to Tyr112 which, due to the new conformation of L3, allows for hydrogen bond formation between Tyr112 and three partners on RBD_wt_: the side chain of Glu414, the backbone carbonyl oxygen atom of Pro412, and the side chain of the acidic Asp427, interactions not present in the native CR3022.

#### H1 CDR design

The Tyr residue in position 28 was replaced by Gly. This mutation seems advantageous because the smaller side chain of glycine and the particular conformation of the designed H1 loop allowed the side chain of Tyr369 from the RBD_wt_ to become buried on the mAb surface. The hydroxyl moiety of Tyr369 can also participate in hydrogen bonding with the side chain -NH group of Trp39, a residue which represents another mutation in the Rosetta-designed H1 loop: Y39W. Trp40 was replaced by a more hydrophobic Phe residue, whose side- chain inserts between the cycle of Tyr380 and the long hydrocarbon chain of Lys387, thus hydrophobically stabilizing the Ab-Ag interaction.

#### H3 CDR design

Serine in position 133 was mutated to a His residue which inserts in between G381 and Asp427 on the RBD_wt_, forming a hydrogen bond-mediated bridge between the -NH group of G381 and the backbone -C=O group of Asp427.

### Physical insights to design a more efficient monoclonal antibody for the RBD

Individual residue electrostatic contributions to the complexation SARS-CoV-2 RBD_wt_-P1 were investigated through the TAS procedure. Each new protein produced with the ALA single mutation (i.e. binder’) is referred to here as P1’. All P1 ionizable residues were replaced by ALA one per time to give P1’, and the corresponding *βw(r)* functions for the complexation SARS- CoV-2 RBD_wt_-P1’ were estimated by the FORTE approach. The differences in the *βw_min_* values observed for P1 and P1’ [ΔΔG=*βw_min_*(P1’)-*βw_min_*(P1)] are portrayed in Figure 9. The purples lines indicate the maximum estimated errors (0.01K_B_T). Free energy shifts larger than 0.01K_B_T are considered significant. Positive shifts indicate mutations that negatively impact the complexation process. These amino acids (ASP-9.LC, GLU-17.LC, ASP-72.LC, ASP-88.LC, ASP-100.LC, GLU-11.HC, GLU-23.HC, GLU-67.HC, ASP-83.HC, and ASP-137.HC) are critical for this molecular mechanism, i.e., replacing one of them with a neutral residue reduces the RBD_wt_ affinity which is not good to block the virus-cell interaction. For instance, the substitution of the ASP residue in position 88 at the light chain (ASP-88.LC) by ALA decreases the binding affinity by 0.03 K_B_T units. Conversely, negative values of ΔΔG correspond to mutations that can improve the capability of P1 to bind the RBD_wt_ which is potentially useful to prevent the viral entry into the host cell. Note that most of the key amino acids that favor the RBD_wt_-P1 complexation are acid residues (ASP and GLU). There are just a couple of exceptions (e.g. ARG-77 at the light chain) but always with smaller differences than the estimated standard deviations (ΔΔG<0.01 K_B_T). The neutralization of a basic ionizable group (e.g. LYS-13 at the heavy chain) implies a reduction of the total net charge of the binder and an improvement on its binding features (ΔΔG_LYS-13.HC_=-0.03 K_B_T). Indeed, it was noticed before that a decrease of the binder net charge gives a higher RBD affinity (Corrêa Giron et al., 2020). This information is a useful insight to design a more efficient therapeutic binder that can avoid the attachment of the RBD to the human cell.

**Figure 9:**
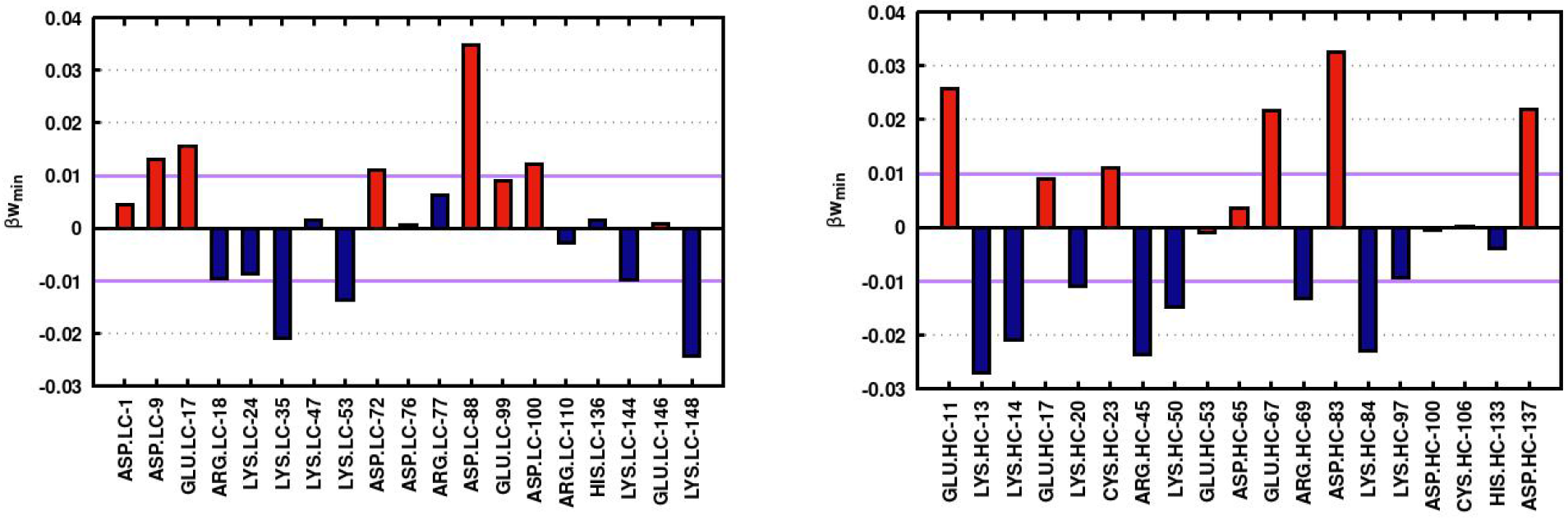
Individual residue electrostatic contributions to the complexation SARS-CoV-2 RBD_wt_-P1. Acid and basic residues are represented by red and blue, respectively. Data from the theoretical ALA scanning procedure applied to P1. Data for the amino acids belonging to the light (LC) and heavy (HC) chains are given in the left and right panels, respectively. Calculations were done with the FORTE approach. ΔΔG (in K_B_T units) is defined as the difference between the minimum value measured in*βw(r)* for the complexation RBD_wt_-P1’ [*β*w_min_(P1’)] and the corresponding quantity for the original P1 [*β*w_min_(P1)] for each new mutation. See the text for more details.

The most important cases where the single ALA mutation in P1 has a stronger influence in the SARS-CoV-2 RBD_wt_-P1 complexation were mapped in the molecular structure. This can be seen in Figure 10.Interestingly not only amino acids at the antigen-antibody interface are critical for the complexation. In fact, amino acids more buried inside the protein structure can also have an important influence on the complexation due to the long-range nature of the electrostatic interactions and the electrostatic coupling between the titratable groups. This is in line with a broader view of epitopes (and paratopes) defined as “electrostatic epitopes” (Poveda-Cuevas et al., 2020), where any ionizable group can affect the interactions and drive the complexation.

**Figure 10:**
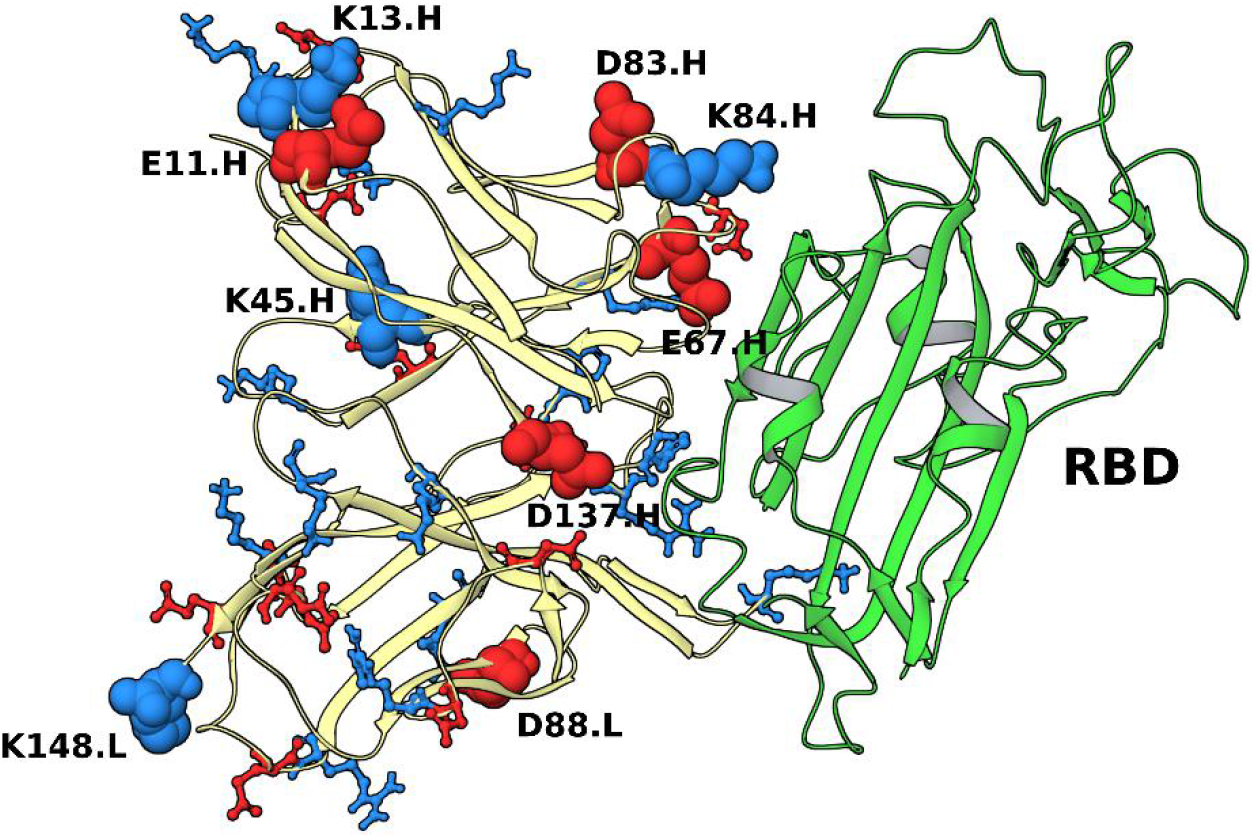
Molecular structure of the SARS-CoV-2 RBD_wt_ complexed with P1. The RBD_wt_ is represented in green. All ionizable residues from the binder P1 whose single ALA mutation resulted in representative ΔΔGs (values larger than the estimated errors) are highlighted by blue (basic residues) and red (acid residues). The most important mutations are labeled. The letters L and H are used to refer to the light and heavy chain, respectively.

### The total net charge of the mAbs and their binding affinities

As mentioned above and previously indicated (Corrêa Giron et al., 2020), there is a tendency for a lower charged mAb to have a higher RBD affinity. In Figure 11a, this correlation is explored for P0 to P10. Although there is no perfect linear behavior, it can be seen that there is indeed this tendency. This is more clearly seen in Figure 11b, where the data from all single, double and triple mutations investigated by the theoretical ALA/GLU scanning is shown together. This data suggests that Coulombic forces are mainly responsible for driving the association of this antigen with the studied mAbs. As a matter of fact, there are now accumulative data on the importance of electrostatic interactions for this system (Bai and Warshel, 2020; Corrêa Giron et al., 2020; Nguyen et al., 2021, 2020). Certainly, other physical interactions can participate in this associative mechanism that is not solely but largely controlled by Coulombic forces. Therefore, the total net charge of the mAb is an important physical-chemical parameter to design better binders with higher affinity. This is what was next explored towards a more specific and efficient macromolecule capable of preventing viral infection.

**Figure 11:**
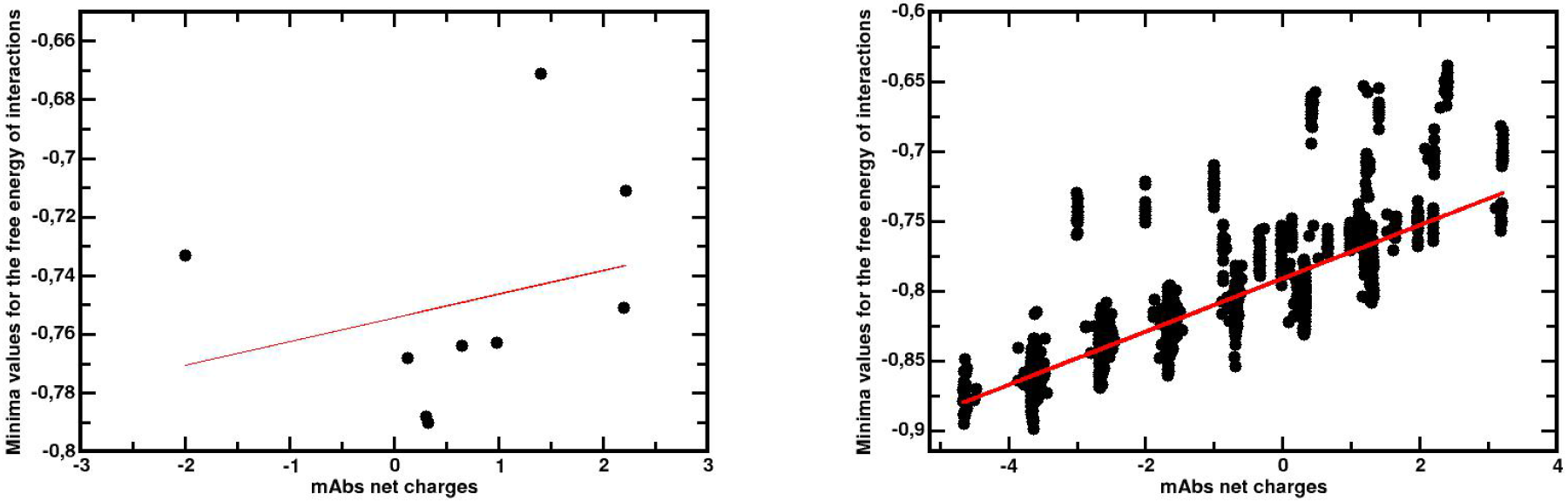
Correlation between the total net charge number of each fragment of mAb and the corresponding *βw_min_*values for their complexations with SARS-CoV-2 RBD_wt_. (a) *Left panel*: Data for each fragment of Rosetta-designed mAb (P1 to P10). (a) *Right panel*: Data for each mutated fragment of P1 and P6 obtained during the theoretical ALA scanning procedure. The data for the total net charge numbers was obtained from titration simulations with a single protein in the absence of the RBD_wt_ by the FPTS at pH 7. All *βw_min_* values were computed from RBD_wt_-complexation studies using FORTE at pH 7 and 150mM of NaCl.

### Additional optimization of the best RBD binder

The two best candidates P1’ (LC: R18E and HC: K20E and K84A) and P6’ (HC: K20A, G61E, and K84A) obtained through the three cycles electrostatic optimization pipeline (based on the TAS procedure) were tested against the most common SARS-CoV-2 mutations. This was done to assess if these macromolecules could have the potential to stop the virus from working efficiently targeting RBDs from common variants. Figures 12a and b show the *βw_min_* values for the complexes RBD-P1’ and RBD-P6’, respectively. The new optimized binders P1’ and P6’ are able to form stable molecular complexes with all studied RBDs. Due to the additional electrostatic optimization, they have improved RBD binding affinities in comparison with the initial template (P01 or P06) provided by the RAbD approach. Consequently, they have more chances for a successful neutralization *in vitro/in vivo* of the virus. Moreover, this result indicated that the integrated biophysical modeling multi-approach used here offers an efficient route to design better macromolecular ligands. Intriguingly both putative mAbs P1’ and P6’ have a stronger affinity for some RBDs from variants that are threatening the immune response. For instance, P1’ has the highest affinity observed for both RBD_Delta_and RBD_Omicron_.This is a promising result to deal with the present crisis offering two ideal candidates to treat the pathogenic threats from different SARS-CoV-2 variants. Quite recently, a pre-print work suggested cross-reactivity specifically between antibodies for Delta and Omicron variants(Khan et al., 2021). P1’ is slightly better than P6’ due to its higher RBD affinity for all mutants in general (e.g. *βw_min_*=-0.96(1)K_B_T for RBD_Delta_-P1’, and *βw_min_*=-0.92(1)K_B_T for RBD_Delta_-P6’). From a bioinformatics point of view, it is interesting to point out that an optimized mAb solely obtained by the three cycles electrostatic optimization pipeline (CR3022’) (Corrêa Giron et al., 2020) is not as good as the one achieved by the present strategy.

**Figure 12:**
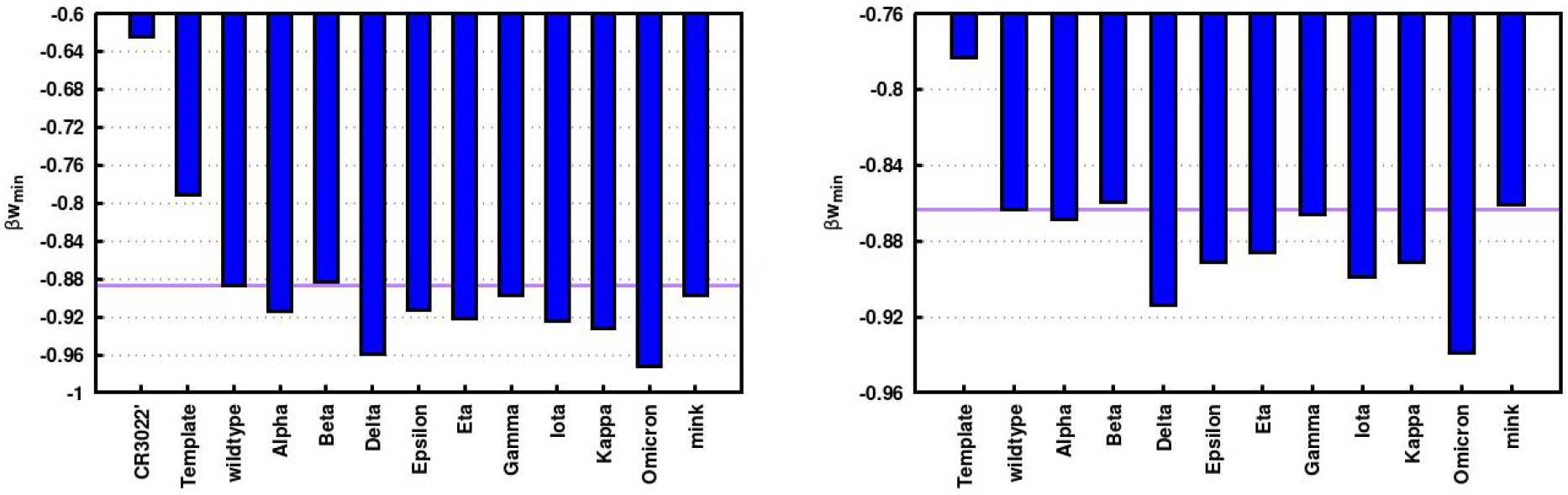
Binding RBD affinities for optimized binders obtained through the three cycles electrostatic optimization pipeline. *(a) Left panel:* Binder based on the Rosetta-designed P1. *(b) Right panel:* Binder based on the Rosetta-designed P2. Data from the estimated *βw_min_*values for the molecular complexation between RBDs from different variants with P1’ (P1 with three mutations LC: R18E and HC: K20E and K84A) and P6’ (P6 with three mutations HC: K20A, G61E, and K84A), respectively. The purple lines are drawn to guide the eyes for the comparison with the outcomes for the RBD_wt_. See the text for more details.

## CONCLUSIONS

RosettaAntibodyDesign protocol was applied to sample the sequence, structure, and binding space of a large number of antibody-antigen complexes. Of totally 235,000 candidates those showing a higher binding affinity compared to the native CR3022-RBD_wt_ complex were identified and finally, 10 complexes were chosen to be evaluated further in binding free energy calculations. Both Umbrella sampling with SIRAH 2.0 coarse-grained force field and the in-house Fast cOarse-grainedpRotein-proTeinmodEl (FORTE) based on fast proton titration scheme (FPTS) were utilized both giving mutually consistent results. After compiling the results at similar conditions to experiments in a heat map we can observe that all the selected RAbD produced CR3022-RBDcomplexes show a stronger binding compared to native CR3022 and also a higher potential to block the interaction between the virus and the host cell. Further, two candidates, P01 and P06, show a clearly strongest interaction.

In a closer analysis, we can find that amino acids both at the surface and also deeper inside the protein structure are critical for the complexation through electrostatic coupling between titratable groups and are characterized as “electrostatic epitopes”.

Furthermore, we can observe that the net charge of the mAbs is one important determinant in binding affinity and there is a clear nearly linear tendency for lower charged mAbs to exhibit higher affinity to RBD. The Coulombic forces are driving the antigen-antibody association, also observed in previous studies (Corrêa Giron et al., 2020; Nguyen et al., 2021), while other short-range interactions are important in the close-range association as well.

We have performed atomistic simulations for the best candidates to study the structural determinants behind the strong affinity of P01 to RBD. The mutations in P01 were in all six complementarity-determining regions (CDR) sequences on both light and heavy chains, improving both the specificity and affinity in favoring the formation of supplementary salt-bridges and H-bonds, as well as through hydrophobic contacts. The two best candidates P01 and P06 were tested against to most common SARS-CoV-2 mutations to hinder the virus targeting RBDs. We could see they both can form stable complexes with all RBDs included in our study thereby having the potential to neutralize the virus both in vitro and in vivo.

Our combined multi-scale approach is found to be a fast, robust, and reliable tool to design better macromolecular ligands allowing us to identify the best candidates for the different variants of SARS-CoV-2 including Omicron. It is a pragmatic approach for a short development cycle for SARS-CoV-2 diagnosis, treatment, and prevention.

This multi-approach is a general theoretical framework towards high specific antibodies for SARS-CoV-2 and can be extended to other diseases (e.g. other viral and bacterial pathogens, leukemia, cancer, multiple sclerosis, rheumatoid arthritis, lupus) both for diagnosis and therapeutic purposes.

## ACKNOWLEDGEMENTS

This work has been supported in part by the “Fundação de Amparo à Pesquisa do Estado de São Paulo” [Fapesp 2020/07158-2 (F.L.B.d.S.)] and the Conselho Nacional de Desenvolvimento Científico e Tecnológico (CNPq) [CNPq 305393/2020-0 (FLBdS)]. F.L.B.d.S. is also deeply thankful for resources provided by the Swedish National Infrastructure for Computing (SNIC) at NSC. A. Laaksonen acknowledges the Swedish Research Council for financial support, and partial support from a grant from the Ministry of Research and Innovation of Romania (CNCS - UEFISCDI, project number PN-III-P4-ID-PCCF-2016-0050, within PNCDI III). F. M. acknowledges financial support from Progetto Fondazione di Sardegna (Grant CUP: F72F20000230007).

## REFERENCES

1. Abraham, M.J., Murtola, T., Schulz, R., Páll, S., Smith, J.C., Hess, B., Lindahl, E., 2015. GROMACS: High performance molecular simulations through multi-level parallelism from laptops to supercomputers. SoftwareX 1–2, 19–25. https://doi.org/10.1016/j.softx.2015.06.001

2. Abraham, M.J., van der Spoel, D., Lindahl, E., Hess, B., and the GROMACS development team, 2019. GROMACS User Manual version 2019. URL http://www.gromacs.org

3. Adolf-Bryfogle, J., Kalyuzhniy, O., Kubitz, M., Weitzner, B.D., Hu, X., Adachi, Y., Schief, W.R., Dunbrack, R.L., 2018. RosettaAntibodyDesign (RAbD): A general framework for computational antibody design. PLOS Comput. Biol. 14, e1006112. https://doi.org/10.1371/journal.pcbi.1006112

4. Adžić, N., Podgornik, R., 2015. Charge regulation in ionic solutions: Thermal fluctuations and Kirkwood-Schumaker interactions. Phys. Rev. E 91. https://doi.org/10.1103/physreve.91.022715

5. Ahmad, B., Batool, M., Kim, M.-S., Choi, S., 2021. Computational-Driven Epitope Verification and Affinity Maturation of TLR4-Targeting Antibodies. Int. J. Mol. Sci. 22, 5989. https://doi.org/10.3390/ijms22115989

6. Alder, B.J., Wainwright, T.E., 1959. Studies in Molecular Dynamics. I. General method. J Chem Phys 31, 459–466.

7. Andreatta, M., Nielsen, M., 2018. Bioinformatics Tools for the Prediction of T-Cell Epitopes, in: Rockberg, J., Nilvebrant, J. (Eds.), Epitope Mapping Protocols, Methods in Molecular Biology. Springer New York, New York, NY, pp. 269–281. https://doi.org/10.1007/978-1-4939-7841-0_18

8. Annavajhala, M.K., Mohri, H., Wang, P., Nair, M., Zucker, J.E., Sheng, Z., Gomez-Simmonds, A., Kelley, A.L., Tagliavia, M., Huang, Y., Bedford, T., Ho, D.D., Uhlemann, A.-C., 2021. Emergence and Expansion of the SARS-CoV-2 Variant B.1.526 Identified in New York. medRxiv 2021.02.23.21252259. https://doi.org/10.1101/2021.02.23.21252259

9. Bai, C., Warshel, A., 2020. Critical Differences between the Binding Features of the Spike Proteins of SARS-CoV-2 and SARS-CoV. J. Phys. Chem. B 124, 5907–5912. https://doi.org/10.1021/acs.jpcb.0c04317

10. Barnes, C.O., Jette, C.A., Abernathy, M.E., Dam, K.-M.A., Esswein, S.R., Gristick, H.B., Malyutin, A.G., Sharaf, N.G., Huey-Tubman, K.E., Lee, Y.E., Robbiani, D.F., Nussenzweig, M.C., West, A.P., Bjorkman, P.J., 2020. SARS-CoV-2 neutralizing antibody structures inform therapeutic strategies. Nature 588, 682–687. https://doi.org/10.1038/s41586-020-2852-1

11. Barroso da Silva, F.L.B. da, MacKernan, D., 2017. Benchmarking a Fast Proton Titration Scheme in Implicit Solvent for Biomolecular Simulations. J Chem Theory Comput 13, 2915–2929.

12. Barroso da Silva, F.L., Carloni, P., Cheung, D., Cottone, G., Donnini, S., Foegeding, E.A., Gulzar, M., Jacquier, J.C., Lobaskin, V., MacKernan, D., Mohammad Hosseini Naveh, Z., Radhakrishnan, R., Santiso, E.E., 2020. Understanding and Controlling Food Protein Structure and Function in Foods: Perspectives from Experiments and Computer Simulations. Annu. Rev. Food Sci. Technol. 11, 365–387. https://doi.org/10.1146/annurev-food-032519-051640

13. Barroso da Silva, F.L., Derreumaux, P., Pasquali, fn, 2018. Protein-RNA complexation driven by the charge regulation mechanism. Biochem. Biophys. Res. Commun., Multiscale Modeling 498, 264–273. https://doi.org/10.1016/j.bbrc.2017.07.027

14. Barroso da Silva, F.L., Dias, L.G., 2017. Development of constant-pH simulation methods in implicit solvent and applications in biomolecular systems. Biophys. Rev. 9, 699–728. https://doi.org/10.1007/s12551-017-0311-5

15. Barroso Da Silva, F.L., Jönsson, B., 2009. Polyelectrolyte-protein complexation driven by charge regulation. Soft Matter 5, 2862–2868.

16. Barroso da Silva, F.L., Lund, M., Jönsson, B., T. \AAkesson, 2006. On the interaction between protein and polyelectrolyte. J Phys Chem B 110, 4459–4464.

17. Barroso da Silva, F.L., Pasquali, S., Derreumaux, P., Dias, L.G., 2016. Electrostatics analysis of the mutational and pH effects of the N-terminal domain self-association of the Major Ampullate Spidroin. Soft Matter 12, 5600–5612.

18. Boggiano, C., Eisinger, R.W., Lerner, A.M., Anderson, J.M., Woodcock, J., Fauci, A.S., Collins, F.S., 2021. Update on and Future Directions for Use of Anti–SARS-CoV-2 Antibodies: National Institutes of Health Summit on Treatment and Prevention of COVID-19. Ann. Intern. Med. M21–3669. https://doi.org/10.7326/M21-3669

19. Boulard, Y., Bressanelli, S., 2021. Recapitulating Trafficking of Nucleosides Into the Active Site of Polymerases of RNA Viruses: The Challenge and the Prize. Front. Med. Technol. 3, 75. https://doi.org/10.3389/fmedt.2021.705875

20. Brielle, E.S., Schneidman-Duhovny, D., Linial, M., 2020. The SARS-CoV-2 Exerts a Distinctive Strategy for Interacting with the ACE2 Human Receptor. Viruses 12, 497. https://doi.org/10.3390/v12050497

21. Burley, S., Berman, H., Kleywegt, G., Markley, J., Nakamura, H., Velankar, S., 2017. Protein Data Bank (PDB): The Single Global Macromolecular Structure Archive, in: Wlodawer, A., Dauter, Z., Jaskolski, M. (Eds.), Protein Crystallography, Methods in Molecular Biology. Springer New York, pp. 627–641. https://doi.org/10.1007/978-1-4939-7000-1_26

22. Callaway, E., 2021. Multitude of coronavirus variants found in the US — but the threat is unclear. Nature 591, 190–190. https://doi.org/10.1038/d41586-021-00564-4

23. Cao, Y., Wang, J., Jian, F., Xiao, T., Song, W., Yisimayi, A., Huang, W., Li, Q., Wang, P., An, R., Wang, J., Wang, Yao, Niu, X., Yang, S., Liang, H., Sun, H., Li, T., Yu, Y., Cui, Q., Liu, S., Yang, X., Du, S., Zhang, Z., Hao, X., Shao, F., Jin, R., Wang, X., Xiao, J., Wang, Youchun, Xie, X.S., 2021. Omicron escapes the majority of existing SARS-CoV-2 neutralizing antibodies. https://doi.org/10.1101/2021.12.07.470392

24. Chen, R.E., Zhang, X., Case, J.B., Winkler, E.S., Liu, Y., VanBlargan, L.A., Liu, J., Errico, J.M., Xie, X., Suryadevara, N., Gilchuk, P., Zost, S.J., Tahan, S., Droit, L., Turner, J.S., Kim, W., Schmitz, A.J., Thapa, M., Wang, D., Boon, A.C.M., Presti, R.M., O’Halloran, J.A., Kim, A.H.J., Deepak, P., Pinto, D., Fremont, D.H., Crowe, J.E., Corti, D., Virgin, H.W., Ellebedy, A.H., Shi, P.-Y., Diamond, M.S., 2021. Resistance of SARS-CoV-2 variants to neutralization by monoclonal and serum-derived polyclonal antibodies. Nat. Med. 27, 717–726. https://doi.org/10.1038/s41591-021-01294-w

25. Chowdhury, R., Allan, M.F., Maranas, C.D., 2018. OptMAVEn-2.0: De novo Design of Variable Antibody Regions Against Targeted Antigen Epitopes. Antibodies 7, 23. https://doi.org/10.3390/antib7030023

26. Chowdhury, R., Grisewood, M.J., Boorla, V.S., Yan, Q., Pfleger, B.F., Maranas, C.D., 2020. IPRO+/−: Computational Protein Design Tool Allowing for Insertions and Deletions. Structure 28, 1344–1357.e4. https://doi.org/10.1016/j.str.2020.08.003

27. Chun Huai Luo, Adannaya Amadi, C. Paul Morris, Matthew Schwartz, Elli Y. Klein, Heba H. Mostafa, 2021. SARS-CoV-2 Variants of concern B.1.1.7 and B.1.351 are not associated with higher viral loads. Presented at the World Microbe Forum 2021, Online worldwide.

28. Corrada, D., Colombo, G., 2013. Energetic and Dynamic Aspects of the Affinity Maturation Process: Characterizing Improved Variants from the Bevacizumab Antibody with Molecular Simulations. J. Chem. Inf. Model. 53, 2937–2950. https://doi.org/10.1021/ci400416e

29. Corrêa Giron, C., Laaksonen, A., Barroso da Silva, F.L., 2020. On the interactions of the receptor-binding domain of SARS-CoV-1 and SARS-CoV-2 spike proteins with monoclonal antibodies and the receptor ACE2. Virus Res. 285, 198021. https://doi.org/10.1016/j.virusres.2020.198021

30. Cortese, M., Neufeldt, C.J., 2021. Exploiting a chink in the armor: engineering broadly neutralizing monoclonal antibodies for SARS-like viruses. Signal Transduct. Target. Ther. 6, 232. https://doi.org/10.1038/s41392-021-00661-w

31. Dascalu, A.I., Ardeleanu, R., Neamtu, A., Maier, S.S., Uritu, C.M., Nicolescu, A., Silion, M., Peptanariu, D., Calin, M., Pinteala, M., 2017. Transfection-capable polycationic nanovectors which include PEGylated-cyclodextrin structural units: a new synthesis pathway. J. Mater. Chem. B 5, 7164–7174. https://doi.org/10.1039/C7TB01722G

32. Delboni, L., Barroso da Silva, F.L., 2016. On the complexation of whey proteins. Food Hydrocoll. 55, 89–99.

33. Deng, X., Garcia-Knight, M.A., Khalid, M.M., Servellita, V., Wang, C., Morris, M.K., Sotomayor-González, A., Glasner, D.R., Reyes, K.R., Gliwa, A.S., Reddy, N.P., Sanchez San Martin, C., Federman, S., Cheng, J., Balcerek, J., Taylor, J., Streithorst, J.A., Miller, S., Sreekumar, B., Chen, P.-Y., Schulze-Gahmen, U., Taha, T.Y., Hayashi, J.M., Simoneau, C.R., Kumar, G.R., McMahon, S., Lidsky, P.V., Xiao, Y., Hemarajata, P., Green, N.M., Espinosa, A., Kath, C., Haw, M., Bell, J., Hacker, J.K., Hanson, C., Wadford, D.A., Anaya, C., Ferguson, D., Frankino, P.A., Shivram, H., Lareau, L.F., Wyman, S.K., Ott, M., Andino, R., Chiu, C.Y., 2021. Transmission, infectivity, and neutralization of a spike L452R SARS-CoV-2 variant. Cell 184, 3426–3437.e8. https://doi.org/10.1016/j.cell.2021.04.025

34. Ding, H., Yin, Y., Ni, S., Sheng, Y., Ma, Y., 2021. Accurate Evaluation on the Interactions of SARS-CoV-2 with Its Receptor ACE2 and Antibodies CR3022/CB6*. Chin. Phys. Lett. 38, 018701. https://doi.org/10.1088/0256-307X/38/1/018701

35. Edara, V.V., Hudson, W.H., Xie, X., Ahmed, R., Suthar, M.S., 2021. Neutralizing Antibodies Against SARS-CoV-2 Variants After Infection and Vaccination. JAMA 325, 1896–1898. https://doi.org/10.1001/jama.2021.4388

36. Engelbrecht, L. de V., Mocci, F., Wang, Y., Perepelytsya, S., Vasiliu, T., Laaksonen, A., 2022. Molecular Perspective on Solutions and Liquid Mixtures from Modelling and Experiment, in: Bulavin, L., Lebovka, N. (Eds.), Soft Matter Systems for Biomedical Applications, Springer Proceedings in Physics. Springer International Publishing, Cham, pp. 53–84. https://doi.org/10.1007/978-3-030-80924-9_3

37. Foote, J., Eisen, H.N., 1995. Kinetic and affinity limits on antibodies produced during immune responses. Proc. Natl. Acad. Sci. 92, 1254–1256. https://doi.org/10.1073/pnas.92.5.1254

38. Ford, C.T., Machado, D.J., Janies, D.A., 2021. Predictions of the SARS-CoV-2 Omicron Variant (B.1.1.529) Spike Protein Receptor-Binding Domain Structure and Neutralizing Antibody Interactions. https://doi.org/10.1101/2021.12.03.471024

39. Francés-Monerris, A., Hognon, C., Miclot, T., García-Iriepa, C., Iriepa, I., Terenzi, A., Grandemange, S., Barone, G., Marazzi, M., Monari, A., 2020. Molecular Basis of SARS-CoV-2 Infection and Rational Design of Potential Antiviral Agents: Modeling and Simulation Approaches. J. Proteome Res. 19, 4291–4315. https://doi.org/10.1021/acs.jproteome.0c00779

40. Fung, M., Otani, I., Pham, M., Babik, J., 2021. Zoonotic coronavirus epidemics. Ann. Allergy. Asthma. Immunol. 126, 321–337. https://doi.org/10.1016/j.anai.2020.11.021

41. Germain, R.N., Meier-Schellersheim, M., Nita-Lazar, A., Fraser, I.D.C., 2011. Systems Biology in Immunology: A Computational Modeling Perspective. Annu. Rev. Immunol. 29, 527– 585. https://doi.org/10.1146/annurev-immunol-030409-101317

42. Giron, C.C., Laaksonen, A., Barroso da Silva, F.L., 2021. Up State of the SARS-COV-2 Spike Homotrimer Favors an Increased Virulence for New Variants. Front. Med. Technol. 3, 694347. https://doi.org/10.3389/fmedt.2021.694347

43. González-Fernández, Á., Bermúdez Silva, F.J., López-Hoyos, M., Cobaleda, C., Montoliu, L., Del Val, M., Leech, K., 2020. Non-animal-derived monoclonal antibodies are not ready to substitute current hybridoma technology. Nat. Methods 17, 1069–1070. https://doi.org/10.1038/s41592-020-00977-5

44. Gunsteren, W.F. van, Luque, F.J., Timms, D., Torda, A.E., 1994. Molecular Mechanics in Biology: From Structure to Function, Taking Account of Solvation. Annu Rev Biophys Biomol Struct. 23, 847–863.

45. Gupta, R.K., Topol, E.J., 2021. COVID-19 vaccine breakthrough infections. Science 374, 1561– 1562. https://doi.org/10.1126/science.abl8487

46. Hastie, K.M., Li, H., Bedinger, D., Schendel, S.L., Dennison, S.M., Li, K., Rayaprolu, V., Yu, X., Mann, C., Zandonatti, M., Diaz Avalos, R., Zyla, D., Buck, T., Hui, S., Shaffer, K., Hariharan, C., Yin, J., Olmedillas, E., Enriquez, A., Parekh, D., Abraha, M., Feeney, E., Horn, G.Q., CoVIC-DB team, Aldon, Y., Ali, H., Aracic, S., Cobb, R.R., Federman, R.S., Fernandez, J.M., Glanville, J., Green, R., Grigoryan, G., Lujan Hernandez, A.G., Ho, D.D., Huang, K.-Y.A., Ingraham, J., Jiang, W., Kellam, P., Kim, C., Kim, M., Kim, H.M., Kong, C., Krebs, S.J., Lan, F., Lang, G., Lee, S., Leung, C.L., Liu, J., Lu, Y., MacCamy, A., McGuire, A.T., Palser, A.L., Rabbitts, T.H., Rikhtegaran Tehrani, Z., Sajadi, M.M., Sanders, R.W., Sato, A.K., Schweizer, L., Seo, J., Shen, B., Snitselaar, J.L., Stamatatos, L., Tan, Y., Tomic, M.T., van Gils, M.J., Youssef, S., Yu, J., Yuan, T.Z., Zhang, Q., Peters, B., Tomaras, G.D., Germann, T., Saphire, E.O., 2021. Defining variant-resistant epitopes targeted by SARS-CoV-2 antibodies: A global consortium study. Science 374, 472–478. https://doi.org/10.1126/science.abh2315

47. Hollingsworth, S.A., Dror, R.O., 2018. Molecular Dynamics Simulation for All. Neuron 99, 1129–1143. https://doi.org/10.1016/j.neuron.2018.08.011

48. Hub, J.S., de Groot, B.L., van der Spoel, D., 2010. g_wham—A Free Weighted Histogram Analysis Implementation Including Robust Error and Autocorrelation Estimates. J. Chem. Theory Comput. 6, 3713–3720. https://doi.org/10.1021/ct100494z

49. Ibrahim, B., McMahon, D.P., Hufsky, F., Beer, M., Deng, L., Mercier, P.L., Palmarini, M., Thiel, V., Marz, M., 2018. A new era of virus bioinformatics. Virus Res. 251, 86–90. https://doi.org/10.1016/j.virusres.2018.05.009

50. Ismail, A.M., Elfiky, A.A., 2020. SARS-CoV-2 spike behavior in situ: a Cryo-EM images for a better understanding of the COVID-19 pandemic. Signal Transduct. Target. Ther. 5, 252. https://doi.org/10.1038/s41392-020-00365-7

51. Jönsson, B., Svensson, B., 1993. Monte Carlo simulation of ion–protein binding, in: Gunsteren, W.F. van, Weiner, P.K., Wilkinson, A. (Eds.), Computer Simulation of Biomolecular Systems. ESCOM, Leiden, pp. 464–482.

52. Jorgensen, W.L., Chandrasekhar, J., Madura, J.D., Impey, R.W., Klein, M.L., 1983. Comparison of simple potential functions for simulating liquid water. J Chem Phys 79, 926–953.

53. Julian, M.C., Li, L., Garde, S., Wilen, R., Tessier, P.M., 2017. Efficient affinity maturation of antibody variable domains requires co-selection of compensatory mutations to maintain thermodynamic stability. Sci. Rep. 7, 45259. https://doi.org/10.1038/srep45259

54. Karplus, M., McCammon, J.A., 2002. Molecular dynamics simulations of biomolecules. Nat. Struc Biol 9, 646–652.

55. Keller, M.A., Stiehm, E.R., 2000. Passive Immunity in Prevention and Treatment of Infectious Diseases. Clin. Microbiol. Rev. 13, 602–614. https://doi.org/10.1128/CMR.13.4.602

56. Keretsu, S., Bhujbal, S.P., Cho, S.J., 2020. Rational approach toward COVID-19 main protease inhibitors via molecular docking, molecular dynamics simulation and free energy calculation. Sci. Rep. 10, 17716. https://doi.org/10.1038/s41598-020-74468-0

57. Khan, K., Karim, F., Cele, S., San, J.E., Lustig, G., Tegally, H., Bernstein, M., Ganga, Y., Jule, Z., Reedoy, K., Ngcobo, N., Mazibuko, M., Mthabela, N., Mhlane, Z., Mbatha, N., Giandhari, J., Ramphal, Y., Naidoo, T., Manickchund, N., Magula, N., Abdool Karim, S.S., Gray, G., Hanekom, W., von Gottberg, A., COMMIT-KZN Team, Gosnell, B.I., Lessells, R.J., Moore, P.L., de Oliveira, T., Moosa, M.-Y.S., Sigal, A., 2021. Omicron infection enhances neutralizing immunity against the Delta variant (preprint). Infectious Diseases (except HIV/AIDS). https://doi.org/10.1101/2021.12.27.21268439

58. Khateeb, J., Li, Y., Zhang, H., 2021. Emerging SARS-CoV-2 variants of concern and potential intervention approaches. Crit. Care 25, 244. https://doi.org/10.1186/s13054-021-03662-x

59. Kim, H.-Y., Stojadinovic, A., Izadjoo, M.J., 2014. Affinity Maturation of Monoclonal Antibodies by Multi-Site-Directed Mutagenesis, in: Ossipow, V., Fischer, N. (Eds.), Monoclonal Antibodies, Methods in Molecular Biology. Humana Press, Totowa, NJ, pp. 407–420. https://doi.org/10.1007/978-1-62703-992-5_24

60. Köhler, G., Milstein, C., 1975. Continuous cultures of fused cells secreting antibody of predefined specificity. Nature 256, 495–497. https://doi.org/10.1038/256495a0

61. Kumar, S., Chandele, A., Sharma, A., 2021. Current status of therapeutic monoclonal antibodies against SARS-CoV-2. PLOS Pathog. 17, e1009885. https://doi.org/10.1371/journal.ppat.1009885

62. Kumar, S., Nussinov, R., 2002. Close-Range Electrostatic Interactions in Proteins. ChemBioChem 3, 604–617. https://doi.org/10.1002/1439-7633(20020703)3:7%3C604::aid-cbic604%3E3.0.co;2-x

63. Kurut, A., Persson, B.A., \AAkesson, T., Forsman, J., Lund, M., 2012. Anisotropic Interactions in Protein Mixtures: Self Assembly and Phase Behavior in Aqueous Solution. J Phys Chem Lett 3, 731–734.

64. Lagoumintzis, G., Chasapis, C.T., Alexandris, N., Kouretas, D., Tzartos, S., Eliopoulos, E., Farsalinos, K., Poulas, K., 2021. Nicotinic cholinergic system and COVID-19: In silico identification of interactions between α7 nicotinic acetylcholine receptor and the cryptic epitopes of SARS-Co-V and SARS-CoV-2 Spike glycoproteins. Food Chem. Toxicol. 149, 112009. https://doi.org/10.1016/j.fct.2021.112009

65. Lapidoth, G.D., Baran, D., Pszolla, G.M., Norn, C., Alon, A., Tyka, M.D., Fleishman, S.J., 2015. *AbDesign* : An algorithm for combinatorial backbone design guided by natural conformations and sequences: Combinatorial Backbone Design in Antibodies. Proteins Struct. Funct. Bioinforma. 83, 1385–1406. https://doi.org/10.1002/prot.24779

66. Laustsen, A.H., Greiff, V., Karatt-Vellatt, A., Muyldermans, S., Jenkins, T.P., 2021. Animal Immunization, in Vitro Display Technologies, and Machine Learning for Antibody Discovery. Trends Biotechnol. 39, 1263–1273. https://doi.org/10.1016/j.tibtech.2021.03.003

67. Li, F., 2016. Structure, Function, and Evolution of Coronavirus Spike Proteins. Annu. Rev. Virol. 3, 237–261. https://doi.org/10.1146/annurev-virology-110615-042301

68. Li, L., Zhang, W., Hu, Y., Tong, X., Zheng, S., Yang, J., Kong, Y., Ren, L., Wei, Q., Mei, H., Hu, C., Tao, C., Yang, R., Wang, Jue, Yu, Y., Guo, Y., Wu, X., Xu, Z., Zeng, L., Xiong, N., Chen, L., Wang, Juan, Man, N., Liu, Y., Xu, H., Deng, E., Zhang, X., Li, C., Wang, C., Su, S., Zhang, L., Wang, Jianwei, Wu, Y., Liu, Z., 2020. Effect of Convalescent Plasma Therapy on Time to Clinical Improvement in Patients With Severe and Life-threatening COVID-19: A Randomized Clinical Trial. JAMA 324, 460. https://doi.org/10.1001/jama.2020.10044

69. Li, W., Zhang, C., Sui, J., Kuhn, J.H., Moore, M.J., Luo, S., Wong, S.-K., Huang, I.-C., Xu, K., Vasilieva, N., Murakami, A., He, Y., Marasco, W.A., Guan, Y., Choe, H., Farzan, M., 2005. Receptor and viral determinants of SARS-coronavirus adaptation to human ACE2. EMBO J. 24, 1634–1643. https://doi.org/10.1038/sj.emboj.7600640

70. Lim, A.W.Y., Williams, G.T., Rada, C., Sale, J.E., 2016. Directed evolution of human scFvs in DT40 cells. Protein Eng. Des. Sel. 29, 39–48. https://doi.org/10.1093/protein/gzv058

71. Lindorff-Larsen, K., Piana, S., Palmo, K., Maragakis, P., Klepeis, J.L., Dror, R.O., Shaw, D.E., 2010. Improved side-chain torsion potentials for the Amber ff99SB protein force field. Proteins Struct. Funct. Bioinforma. 78, 1950–1958. https://doi.org/10.1002/prot.22711

72. Lippow, S.M., Wittrup, K.D., Tidor, B., 2007. Computational design of antibody-affinity improvement beyond in vivo maturation. Nat. Biotechnol. 25, 1171–1176. https://doi.org/10.1038/nbt1336

73. Lou, J., Marks, J.D., 2010. Affinity Maturation by Chain Shuffling and Site Directed Mutagenesis, in: Kontermann, R., Dübel, S. (Eds.), Antibody Engineering. Springer Berlin Heidelberg, Berlin, Heidelberg, pp. 377–396. https://doi.org/10.1007/978-3-642-01144-3_25

74. Lund, M., Jönsson, B., 2013. Charge regulation in biomolecular solution. Q. Rev. Biophys. 46, 265–281.

75. Lund, M., Vrbka, L., Jungwirth, P., 2008. Specific Ion Binding to Nonpolar Surface Patches of Proteins. J. Am. Chem. Soc. 130, 11582–11583. https://doi.org/10.1021/ja803274p

76. Lyubartsev, A., Laaksonen, A., 2021. Inverse Problems and Hierarchical Multiscale Modelling of Biological Matter, in: J.M. Abadie, M., Pinteala, M., Rotaru, A. (Eds.), New Trends in Macromolecular and Supramolecular Chemistry for Biological Applications. Springer International Publishing, Cham, pp. 213–237. https://doi.org/10.1007/978-3-030-57456-7_11

77. Machado, M.R., Barrera, E.E., Klein, F., Sóñora, M., Silva, S., Pantano, S., 2019. The SIRAH 2.0 Force Field: Altius, Fortius, Citius. J. Chem. Theory Comput. 15, 2719–2733. https://doi.org/10.1021/acs.jctc.9b00006

78. Maguire, J.B., Haddox, H.K., Strickland, D., Halabiya, S.F., Coventry, B., Griffin, J.R., Pulavarti, S.V.S.R.K., Cummins, M., Thieker, D.F., Klavins, E., Szyperski, T., DiMaio, F., Baker, D., Kuhlman, B., 2021. Perturbing the energy landscape for improved packing during computational protein design. Proteins Struct. Funct. Bioinforma. 89, 436–449. https://doi.org/10.1002/prot.26030

79. Mahajan, S.P., Meksiriporn, B., Waraho-Zhmayev, D., Weyant, K.B., Kocer, I., Butler, D.C., Messer, A., Escobedo, F.A., DeLisa, M.P., 2018. Computational affinity maturation of camelid single-domain intrabodies against the nonamyloid component of alpha-synuclein. Sci. Rep. 8, 17611. https://doi.org/10.1038/s41598-018-35464-7

80. Mallapaty, S., Callaway, E., Kozlov, M., Ledford, H., Pickrell, J., Van Noorden, R., 2021. How COVID vaccines shaped 2021 in eight powerful charts. Nature 600, 580–583. https://doi.org/10.1038/d41586-021-03686-x

81. Marcus, R.A., 1955. Calculation of thermodynamic properties of polyelectrolytes. J Chem Phys 23, 1057.

82. Martí, D., Alsina, M., Alemán, C., Bertran, O., Turon, P., Torras, J., 2021. Unravelling the molecular interactions between the SARS-CoV-2 RBD spike protein and various specific monoclonal antibodies. Biochimie S0300-9084(21)00249–2. https://doi.org/10.1016/j.biochi.2021.10.013

83. McCammon, J.A., Gelin, B.R., Karplus, M., 1977. Dynamics of folded proteins. Nature 267, 585–590. https://doi.org/10.1038/267585a0

84. Mendonça, D.C., Macedo, J.N., Guimarães, S.L., Barroso da Silva, F.L., Cassago, A., Garratt, R.C., Portugal, R.V., Araujo, A.P.U., 2019. A revised order of subunits in mammalian septin complexes. Cytoskeleton 76, 457–466. https://doi.org/10.1002/cm.21569

85. Metropolis, N.A., Rosenbluth, A.W., Rosenbluth, M.N., Teller, A., Teller, E., 1953. Equation of State Calculations by Fast Computing Machines. J Chem Phys 21, 1087–1097.

86. Minea, B., Marangoci, N., Peptanariu, D., Rosca, I., Nastasa, V., Corciova, A., Varganici, C., Nicolescu, A., Fifere, A., Neamtu, A., Mares, M., Barboiu, M., Pinteala, M., 2016. Inclusion complexes of propiconazole nitrate with substituted β-cyclodextrins: the synthesis and in silico and in vitro assessment of their antifungal properties. New J. Chem. 40, 1765–1776. https://doi.org/10.1039/C5NJ01811K

87. Moal, I.H., Fernández-Recio, J., 2012. SKEMPI: a Structural Kinetic and Energetic database of Mutant Protein Interactions and its use in empirical models. Bioinformatics 28, 2600– 2607. https://doi.org/10.1093/bioinformatics/bts489

88. Mondal, D., Warshel, A., 2020. Exploring the Mechanism of Covalent Inhibition: Simulating the Binding Free Energy of α-Ketoamide Inhibitors of the Main Protease of SARS-CoV-2. Biochemistry 59, 4601–4608. https://doi.org/10.1021/acs.biochem.0c00782

89. Moreira, R.A., Guzman, H.V., Boopathi, S., Baker, J.L., Poma, A.B., 2020. Characterization of Structural and Energetic Differences between Conformations of the SARS-CoV-2 Spike Protein. Materials 13, 5362. https://doi.org/10.3390/ma13235362

90. Natarajan, H., Crowley, A.R., Butler, S.E., Xu, S., Weiner, J.A., Bloch, E.M., Littlefield, K., Wieland-Alter, W., Connor, R.I., Wright, P.F., Benner, S.E., Bonny, T.S., Laeyendecker, O., Sullivan, D., Shoham, S., Quinn, T.C., Larman, H.B., Casadevall, A., Pekosz, A., Redd, A.D., Tobian, A.A.R., Ackerman, M.E., 2021. Markers of Polyfunctional SARS-CoV-2 Antibodies in Convalescent Plasma. mBio 12. https://doi.org/10.1128/mBio.00765-21

91. Nguyen, H., Lan, P.D., Nissley, D.A., O’Brien, E.P., Li, M.S., 2021. Electrostatic Interactions Explain the Higher Binding Affinity of the CR3022 Antibody for SARS-CoV-2 than the 4A8 Antibody. J. Phys. Chem. B 125, 7368–7379. https://doi.org/10.1021/acs.jpcb.1c03639

92. Nguyen, H.L., Lan, P.D., Thai, N.Q., Nissley, D.A., O’Brien, E.P., Li, M.S., 2020. Does SARS-CoV-2 Bind to Human ACE2 More Strongly Than Does SARS-CoV? J. Phys. Chem. B 124, 7336–7347. https://doi.org/10.1021/acs.jpcb.0c04511

93. Noy-Porat, T., Makdasi, E., Alcalay, R., Mechaly, A., Levy, Y., Bercovich-Kinori, A., Zauberman, A., Tamir, H., Yahalom-Ronen, Y., Israeli, M., Epstein, E., Achdout, H., Melamed, S., Chitlaru, T., Weiss, S., Peretz, E., Rosen, O., Paran, N., Yitzhaki, S., Shapira, S.C., Israely, T., Mazor, O., Rosenfeld, R., 2020. A panel of human neutralizing mAbs targeting SARS-CoV-2 spike at multiple epitopes. Nat. Commun. 11, 4303. https://doi.org/10.1038/s41467-020-18159-4

94. Ó Conchúir, S., Barlow, K.A., Pache, R.A., Ollikainen, N., Kundert, K., O’Meara, M.J., Smith, C.A., Kortemme, T., 2015. A Web Resource for Standardized Benchmark Datasets, Metrics, and Rosetta Protocols for Macromolecular Modeling and Design. PLOS ONE 10, e0130433. https://doi.org/10.1371/journal.pone.0130433

95. Pantazes, R.J., Maranas, C.D., 2010. OptCDR: a general computational method for the design of antibody complementarity determining regions for targeted epitope binding. Protein Eng. Des. Sel. 23, 849–858. https://doi.org/10.1093/protein/gzq061

96. Pappas, N., Roux, S., Hölzer, M., Lamkiewicz, K., Mock, F., Marz, M., Dutilh, B.E., 2021. Virus Bioinformatics, in: Encyclopedia of Virology. Elsevier, pp. 124–132. https://doi.org/10.1016/B978-0-12-814515-9.00034-5

97. Parray, H.A., Shukla, S., Samal, S., Shrivastava, T., Ahmed, S., Sharma, C., Kumar, R., 2020. Hybridoma technology a versatile method for isolation of monoclonal antibodies, its applicability across species, limitations, advancement and future perspectives. Int. Immunopharmacol. 85, 106639. https://doi.org/10.1016/j.intimp.2020.106639

98. Patel, J.S., Ytreberg, F.M., 2018. Fast Calculation of Protein–Protein Binding Free Energies Using Umbrella Sampling with a Coarse-Grained Model. J. Chem. Theory Comput. 14, 991–997. https://doi.org/10.1021/acs.jctc.7b00660

99. Persson, B., Lund, M., Forsman, J., Chatterton, D.E.W., Torbjörn \AAkesson, 2010. Molecular evidence of stereo-specific lactoferrin dimers in solution. Biophys Chem 3, 187–189.

100. Persson, H., Kirik, U., Thörnqvist, L., Greiff, L., Levander, F., Ohlin, M., 2018. In Vitro Evolution of Antibodies Inspired by In Vivo Evolution. Front. Immunol. 9, 1391. https://doi.org/10.3389/fimmu.2018.01391

101. Pettersen, E.F., Goddard, T.D., Huang, C.C., Greenblatt, G.S., Meng, E.C., Ferrin, T.E., 2004. UCSF Chimera: a Visualization System for Exploratory Research and Analysis. J Comp Chem 25, 1605–1612.

102. Pons, J., Rajpal, A., Kirsch, J.F., 1999. Energetic analysis of an antigen/antibody interface: Alanine scanning mutagenesis and double mutant cycles on the hyhel-10/lysozyme interaction. Protein Sci. 8, 958–968. https://doi.org/10.1110/ps.8.5.958

103. Poveda-Cuevas, S.A., Barroso da Silva, F.L., Etchebest, C., 2021. How the Strain Origin of Zika Virus NS1 Protein Impacts Its Dynamics and Implications to Their Differential Virulence. J. Chem. Inf. Model. acs.jcim.0c01377. https://doi.org/10.1021/acs.jcim.0c01377

104. Poveda-Cuevas, S.A., Etchebest, C., Barroso da Silva, F.L., 2020. Identification of Electrostatic Epitopes in Flavivirus by Computer Simulations: The PROCEEDpKa Method. J. Chem. Inf. Model. 60, 944–963. https://doi.org/10.1021/acs.jcim.9b00895

105. Poveda-Cuevas, S.A., Etchebest, C., Barroso da Silva, F.L., 2018. Insights into the ZIKV NS1 Virology from Different Strains through a Fine Analysis of Physicochemical Properties. ACS Omega 3, 16212–16229. https://doi.org/10.1021/acsomega.8b02081

106. Prates-Syed, W.A., Chaves, L.C.S., Crema, K.P., Vuitika, L., Lira, A., Côrtes, N., Kersten, V., Guimarães, F.E.G., Sadraeian, M., Barroso da Silva, F.L., Cabral-Marques, O., Barbuto, J.A.M., Russo, M., Câmara, N.O.S., Cabral-Miranda, G., 2021. VLP-Based COVID-19 Vaccines: An Adaptable Technology against the Threat of New Variants. Vaccines 9, 1409. https://doi.org/10.3390/vaccines9121409

107. Pruett, P.S., Air, G.M., 1998. Critical interactions in binding antibody NC41 to influenza N9 neuraminidase: amino acid contacts on the antibody heavy chain. Biochemistry 37, 10660–10670. https://doi.org/10.1021/bi9802059

108. Rahman, A., 1964. Correlations in the motion of atoms in liquid argon. Phys Rev 136, A405– A411.

109. Rapaport, D.C., 2018. Molecular dynamics study of T = 3 capsid assembly. J. Biol. Phys. 44, 147–162. https://doi.org/10.1007/s10867-018-9486-7

110. Reis, G., dos Santos Moreira-Silva, E.A., Silva, D.C.M., Thabane, L., Milagres, A.C., Ferreira, T.S., dos Santos, C.V.Q., de Souza Campos, V.H., Nogueira, A.M.R., de Almeida, A.P.F.G., Callegari, E.D., de Figueiredo Neto, A.D., Savassi, L.C.M., Simplicio, M.I.C., Ribeiro, L.B., Oliveira, R., Harari, O., Forrest, J.I., Ruton, H., Sprague, S., McKay, P., Glushchenko, A.V., Rayner, C.R., Lenze, E.J., Reiersen, A.M., Guyatt, G.H., Mills, E.J., 2022. Effect of early treatment with fluvoxamine on risk of emergency care and hospitalisation among patients with COVID-19: the TOGETHER randomised, platform clinical trial. Lancet Glob. Health 10, e42–e51. https://doi.org/10.1016/S2214-109X(21)00448-4

111. Riahi, S., Lee, J.H., Wei, S., Cost, R., Masiero, A., Prades, C., Olfati-Saber, R., Wendt, M., Park, A., Qiu, Y., Zhou, Y., 2021. Application of an integrated computational antibody engineering platform to design SARS-CoV-2 neutralizers. Antib. Ther. tbab011. https://doi.org/10.1093/abt/tbab011

112. Rubin, R., 2021. Monoclonal Antibodies for COVID-19 Preexposure Prophylaxis Can’t Come Fast Enough for Some People. JAMA 326, 1895. https://doi.org/10.1001/jama.2021.19534

113. Salazar, E., Kuchipudi, S.V., Christensen, P.A., Eagar, T.N., Yi, X., Zhao, P., Jin, Z., Long, S.W., Olsen, R.J., Chen, J., Castillo, B., Leveque, C., Towers, D.M., Lavinder, J., Gollihar, J.D., Cardona, J., Ippolito, G.C., Nissly, R.H., Bird, I.M., Greenawalt, D., Rossi, R.M., Gontu, A., Srinivasan, S., Poojary, I.B., Cattadori, I.M., Hudson, P.J., Joselyn, N., Prugar, L., Huie, K., Herbert, A., Bernard, D.W., Dye, J., Kapur, V., Musser, J.M., 2020. Relationship between Anti-Spike Protein Antibody Titers and SARS-CoV-2 *In Vitro* Virus Neutralization in Convalescent Plasma (preprint). Immunology. https://doi.org/10.1101/2020.06.08.138990

114. Sato, H., Yokoyama, M., Toh, H., 2013. Genomics and computational science for virus research. Front. Microbiol. 4. https://doi.org/10.3389/fmicb.2013.00042

115. Schlick, T., Portillo-Ledesma, S., 2021. Biomolecular modeling thrives in the age of technology. Nat. Comput. Sci. 1, 321–331. https://doi.org/10.1038/s43588-021-00060-9

116. Shariatifar, H., Farasat, A., 2021. Affinity enhancement of CR3022 binding to RBD; in silico site directed mutagenesis using molecular dynamics simulation approaches. J. Biomol. Struct. Dyn. 0, 1–10. https://doi.org/10.1080/07391102.2021.2004230

117. Sharma, D., Priyadarshini, P., Vrati, S., 2015. Unraveling the Web of Viroinformatics: Computational Tools and Databases in Virus Research. J. Virol. 89, 1489–1501. https://doi.org/10.1128/JVI.02027-14

118. Sironi, M., Kaderali, L., 2021. Bioinformatics Algorithms and Predictive Models: The Grand Challenge in Computational Virology. Front. Virol. 1, 684608. https://doi.org/10.3389/fviro.2021.684608

119. Sivasubramanian, A., Sircar, A., Chaudhury, S., Gray, J.J., 2009. Toward high-resolution homology modeling of antibody F v regions and application to antibody-antigen docking. Proteins Struct. Funct. Bioinforma. 74, 497–514. https://doi.org/10.1002/prot.22309

120. Steinhauser, M., Hiermaier, S., 2009. A Review of Computational Methods in Materials Science: Examples from Shock-Wave and Polymer Physics. Int. J. Mol. Sci. 10, 5135–5216. https://doi.org/10.3390/ijms10125135

121. Svensson, B., Jönsson, B., Woodward, C.E., 1990. Electrostatic Contributions of the Binding of Ca^2+$^ in Calbindin mutants. A Monte Carlo Study. Biophys Chem 38, 179–183.

122. Tachioka, M., Sugimoto, N., Nakamura, A., Sunagawa, N., Ishida, T., Uchiyama, T., Igarashi, K., Samejima, M., 2016. Development of simple random mutagenesis protocol for the protein expression system in Pichia pastoris. Biotechnol. Biofuels 9, 199. https://doi.org/10.1186/s13068-016-0613-z

123. Teixeira, A.A., Lund, M., Silva, F.L.B., 2010. Fast Proton Titration Scheme for Multiscale Modeling of Protein Solutions. J. Chem. Theory Comput. 6, 3259–3266.

124. Tian, X., Li, C., Huang, A., Xia, S., Lu, S., Shi, Z., Lu, L., Jiang, S., Yang, Z., Wu, Y., Ying, T., 2020. Potent binding of 2019 novel coronavirus spike protein by a SARS coronavirus-specific human monoclonal antibody. Emerg. Microbes Infect. 9, 382–385. https://doi.org/10.1080/22221751.2020.1729069

125. Tiller, K.E., Chowdhury, R., Li, T., Ludwig, S.D., Sen, S., Maranas, C.D., Tessier, P.M., 2017. Facile Affinity Maturation of Antibody Variable Domains Using Natural Diversity Mutagenesis. Front. Immunol. 8, 986. https://doi.org/10.3389/fimmu.2017.00986

126. Tso, F.Y., Lidenge, S.J., Poppe, L.K., Peña, P.B., Privatt, S.R., Bennett, S.J., Ngowi, J.R., Mwaiselage, J., Belshan, M., Siedlik, J.A., Raine, M.A., Ochoa, J.B., Garcia-Diaz, J., Nossaman, B., Buckner, L., Roberts, W.M., Dean, M.J., Ochoa, A.C., West, J.T., Wood, C., 2021. Presence of antibody-dependent cellular cytotoxicity (ADCC) against SARS-CoV-2 in COVID-19 plasma. PLOS ONE 16, e0247640. https://doi.org/10.1371/journal.pone.0247640

127. van Gunsteren, W.F., Dolenc, J., 2012. Thirty-five years of biomolecular simulation: development of methodology, force fields and software. Mol. Simul. 38, 1271–1281. https://doi.org/10.1080/08927022.2012.701744

128. van Zundert, G.C.P., Rodrigues, Trellet, M., Schmitz, C., Kastritis, P.L., Karaca, E., Melquiond, A.S.J., van Dijk, M., de Vries, S.J., Bonvin, A.M.J.J., 2016. The HADDOCK2.2 Web Server: User-Friendly Integrative Modeling of Biomolecular Complexes. J. Mol. Biol. 428, 720–725. https://doi.org/10.1016/j.jmb.2015.09.014

129. Verkhivker, G.M., Di Paola, L., 2021. Integrated Biophysical Modeling of the SARS-CoV-2 Spike Protein Binding and Allosteric Interactions with Antibodies. J. Phys. Chem. B 125, 4596– 4619. https://doi.org/10.1021/acs.jpcb.1c00395

130. Vilar, S., Cozza, G., Moro, S., 2008. Medicinal Chemistry and the Molecular Operating Environment (MOE): Application of QSAR and Molecular Docking to Drug Discovery. Curr. Top. Med. Chem. 8, 1555–1572. https://doi.org/10.2174/156802608786786624

131. V’kovski, P., Kratzel, A., Steiner, S., Stalder, H., Thiel, V., 2021. Coronavirus biology and replication: implications for SARS-CoV-2. Nat. Rev. Microbiol. 19, 155–170. https://doi.org/10.1038/s41579-020-00468-6

132. Wade, R.C., Gabdoulline, R.G., Lüdemann, S.K., Lounnas, V., 1998. Electrostatic steering and ionic tethering in enzyme-ligand binding: Insights from simulations. Proc Natl Acad Sci USA 95, 5942–5949.

133. Walls, A.C., Park, Y.-J., Tortorici, M.A., Wall, A., McGuire, A.T., Veesler, D., 2020. Structure, Function, and Antigenicity of the SARS-CoV-2 Spike Glycoprotein. Cell S0092867420302622. https://doi.org/10.1016/j.cell.2020.02.058

134. Wang, L., Zhou, T., Zhang, Y., Yang, E.S., Schramm, C.A., Shi, W., Pegu, A., Oloniniyi, O.K., Henry, A.R., Darko, S., Narpala, S.R., Hatcher, C., Martinez, D.R., Tsybovsky, Y., Phung, E., Abiona, O.M., Antia, A., Cale, E.M., Chang, L.A., Choe, M., Corbett, K.S., Davis, R.L., DiPiazza, A.T., Gordon, I.J., Hait, S.H., Hermanus, T., Kgagudi, P., Laboune, F., Leung, K., Liu, T., Mason, R.D., Nazzari, A.F., Novik, L., O’Connell, S., O’Dell, S., Olia, A.S., Schmidt, S.D., Stephens, T., Stringham, C.D., Talana, C.A., Teng, I.-T., Wagner, D.A., Widge, A.T., Zhang, B., Roederer, M., Ledgerwood, J.E., Ruckwardt, T.J., Gaudinski, M.R., Moore, P.L., Doria-Rose, N.A., Baric, R.S., Graham, B.S., McDermott, A.B., Douek, D.C., Kwong, P.D., Mascola, J.R., Sullivan, N.J., Misasi, J., 2021. Ultrapotent antibodies against diverse and highly transmissible SARS-CoV-2 variants. Science 373, eabh1766. https://doi.org/10.1126/science.abh1766

136. Weitzner, B.D., Jeliazkov, J.R., Lyskov, S., Marze, N., Kuroda, D., Frick, R., Adolf-Bryfogle, J., Biswas, N., Dunbrack, R.L., Gray, J.J., 2017. Modeling and docking of antibody structures with Rosetta. Nat. Protoc. 12, 401–416. https://doi.org/10.1038/nprot.2016.180

137. Whittaker, G.R., 2021. SARS-CoV-2 spike and its adaptable furin cleavage site. Lancet Microbe 2, e488–e489. https://doi.org/10.1016/S2666-5247(21)00174-9

138. Wodak, S.J., Janin, J., 2002. Structural basis of macromolecular recognition, in: Advances in Protein Chemistry, Protein Modules and Protein-Protein Interaction. Academic Press, pp. 9–73. https://doi.org/10.1016/S0065-3233(02)61001-0

139. Wu, N.C., Yuan, M., Bangaru, S., Huang, D., Zhu, X., Lee, C.-C.D., Turner, H.L., Peng, L., Yang, L., Burton, D.R., Nemazee, D., Ward, A.B., Wilson, I.A., 2020. A natural mutation between SARS-CoV-2 and SARS-CoV determines neutralization by a cross-reactive antibody. PLOS Pathog. 16, e1009089. https://doi.org/10.1371/journal.ppat.1009089

140. Xie, Y., Guo, W., Lopez-Hernadez, A., Teng, S., Li, L., 2021. The pH Effects on SARS-CoV and SARS-CoV-2 Spike Proteins in the Process of Binding to hACE2. Res. Sq. rs.3.rs-871118. https://doi.org/10.21203/rs.3.rs-871118/v1

141. Yan, R., Zhang, Y., Li, Y., Ye, F., Guo, Y., Xia, L., Zhong, X., Chi, X., Zhou, Q., 2021. Structural basis for the different states of the spike protein of SARS-CoV-2 in complex with ACE2. Cell Res. 31, 717–719. https://doi.org/10.1038/s41422-021-00490-0

142. Yuan, M., Wu, N.C., Zhu, X., Lee, C.-C.D., So, R.T.Y., Lv, H., Mok, C.K.P., Wilson, I.A., 2020. A highly conserved cryptic epitope in the receptor-binding domains of SARS-CoV-2 and SARS-CoV. Science eabb7269. https://doi.org/10.1126/science.abb7269

143. Zaroff, S., Tan, G., 2019. Hybridoma technology: the preferred method for monoclonal antibody generation for *in vivo* applications. BioTechniques 67, 90–92. https://doi.org/10.2144/btn-2019-0054

