## SupplementaryMaterial for "Towards an optimal monoclonal antibody with higher binding affinity to the receptor-binding domain of SARS-CoV-2 spike proteins from different variants"

### Supplementary information

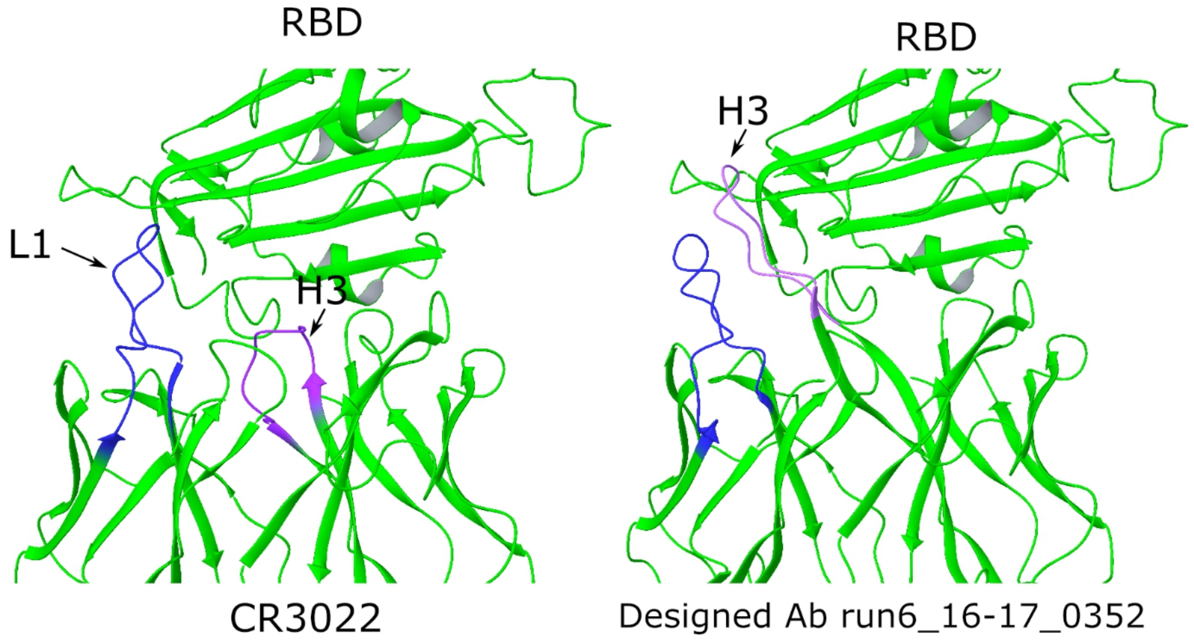


**Figure S1.** Example of a candidate mAb considered false positive. Native CR3022/RBD complex (left) and the RAbD predicted pose, code 16-17_0352, of mAb-RBD complex (right) have been represented with the L1 and H3 CDRs highlighted. It can be seen how the designed long H3 (mauve) CDR actually prevents native L1 (blue) CDR from interacting with the antigen. The contact area between the 16-17_0352 mAb and RBD is higher (2339 Å$^2$) than in the native complex (2060 Å$^2$). However, H3 modelling should be regarded with care, especially for long H3 CDRs, for which the predictions have a low reliability.

**Table S1.** mAb candidates that meet the selection criteria described in the "Theoretical methods" section of the paper main body (ΔG < - 150 REU, SASA > 1900 Å$^2$). Red is for the (A) approach; blue for (B) and black for (C) approaches as described in the main text.

| | ΔG (REU) | SASA h (Å$^2$) | SASA (Å$^2$) | SASAp (Å$^2$) | nH B | nres | Candidate code | |
|---|---|---|---|---|---|---|---|---|
| 1 | -362 | 1078 | 2009 | 932 | 9 | 80 | INTERFACE_rescored_01.pose.6w41-FabOut.10_0119_0001 | P01 |
| 2 | -357 | 1022 | 1979 | 957 | 8 | 78 | INTERFACE_rescored_01.pose.6w41-FabOut.05_0483_0001 | P02 |
| 3 | -298 | 1141 | 1955 | 814 | 3 | 86 | INTERFACE_rescored_02.pose.6w41-FabOut.05_0427_0001 | P03 |
| 4 | -282 | 1058 | 2029 | 971 | 10 | 94 | INTERFACE_rescored_03.pose.6w41-FabOut.24_0219_0001 | P04 |
| 5 | -272 | 1166 | 1941 | 775 | 7 | 80 | INTERFACE_rescored_04.pose.6w41-FabOut.05_0291_0001 | P05 |
| 6 | -251 | 1087 | 1995 | 908 | 6 | 83 | INTERFACE_rescored_03.pose.6w41-FabOut.03_0220_0001 | P06 |
| 7 | -248 | 1190 | 2065 | 875 | 6 | 93 | INTERFACE_rescored_08.pose.6w41-FabOut.08_0886_0001 | P07 |
| 8 | -242 | 1155 | 1994 | 839 | 7 | 81 | INTERFACE_rescored_10.pose.6w41-FabOut.21_00364_0001 | P08 |
| 9 | -230 | 1110 | 1966 | 856 | 8 | 81 | INTERFACE_rescored_18.pose.6w41-FabOut.06_00190_0001 | P09 |
| 10 | -227 | 1092 | 1932 | 840 | 8 | 81 | INTERFACE_rescored_07.pose.6w41-FabOut.03_0550_0001 | P10 |
| 11 | -226 | 1135 | 1919 | 785 | 5 | 82 | INTERFACE_rescored_12.pose.6w41-FabOut.08_0558_0001 | |
| 12 | -220 | 977 | 1916 | 939 | 9 | 86 | INTERFACE_rescored_01.pose.6w41-FabOut.01_0110_0001 | |
| 13 | -201 | 986 | 1928 | 942 | 8 | 81 | INTERFACE_rescored_14.pose.6w41-FabOut.19_0918_0001 | |
| 14 | -198 | 1048 | 1999 | 952 | 9 | 86 | INTERFACE_rescored_11.pose.6w41-FabOut.03_0437_0001 | |
| 15 | -186 | 1156 | 1951 | 795 | 5 | 83 | INTERFACE_rescored_13.pose.6w41-FabOut.24_0566_0001 | |
| 16 | -185 | 1100 | 2005 | 905 | 11 | 84 | INTERFACE_rescored_11.pose.6w41-FabOut.20_0689_0001 | |
| 17 | -184 | 1053 | 1968 | 915 | 7 | 80 | INTERFACE_rescored_17.pose.6w41-FabOut.09_0149_0001 | |
| 18 | -181 | 1102 | 1995 | 893 | 9 | 82 | INTERFACE_rescored_19.pose.6w41-FabOut.02_0806_0001 | |
| 19 | -178 | 1099 | 2082 | 982 | 9 | 85 | INTERFACE_rescored_18.pose.6w41-FabOut.17_0043_0001 | |
| 20 | -177 | 997 | 1954 | 957 | 11 | 78 | INTERFACE_rescored_16.pose.6w41-FabOut.19_0451_0001 | |
| 21 | -177 | 1016 | 1940 | 924 | 9 | 81 | INTERFACE_rescored_13.pose.6w41-FabOut.15_0068_0001 | |
| 22 | -176 | 1115 | 2052 | 937 | 10 | 85 | INTERFACE_rescored_15.pose.6w41-FabOut.04_0788_0001 | |
| 23 | -175 | 1013 | 1907 | 893 | 8 | 79 | INTERFACE_rescored_10.pose.6w41-FabOut.08_0719_0001 | |
| 24 | -175 | 1040 | 1930 | 890 | 9 | 85 | INTERFACE_rescored_14.pose.6w41-FabOut.13_0109_0001 | |
| 25 | -175 | 1184 | 2079 | 895 | 8 | 81 | INTERFACE_rescored_20.pose.6w41-FabOut.02_0069_0001 | |
| 26 | -175 | 1067 | 1937 | 870 | 7 | 83 | INTERFACE_rescored_12.pose.6w41-FabOut.02_0701_0001 | |
| 27 | -175 | 1133 | 2000 | 867 | 8 | 85 | INTERFACE_rescored_17.pose.6w41-FabOut.08_0226_0001 | |
| 28 | -174 | 1117 | 1911 | 794 | 8 | 82 | INTERFACE_rescored_20.pose.6w41-FabOut.23_0912_0001 | |
| 29 | -173 | 1042 | 1959 | 917 | 8 | 83 | INTERFACE_rescored_06.pose.6w41-FabOut.14_0537_0001 | |
| 30 | -172 | 1004 | 1989 | 984 | 10 | 81 | INTERFACE_rescored_14.pose.6w41-FabOut.12_0003_0001 | |
| 31 | -170 | 1072 | 1925 | 853 | 8 | 82 | INTERFACE_rescored_05.pose.6w41-FabOut.18_0099_0001 | |
| 32 | -170 | 959 | 1903 | 944 | 10 | 78 | INTERFACE_rescored_18.pose.6w41-FabOut.16_0459_0001 | |
| 33 | -165 | 1047 | 1968 | 921 | 11 | 83 | INTERFACE_rescored_16.pose.6w41-FabOut.13_0289_0001 | |

| | | | | | | | |
|---|---|---|---|---|---|---|---|
| 34 | -165 | 1105 | 2189 | 1084 | 8 | 80 | INTERFACE_rescored_20.pose.6w41-FabOut.10_0302_0001 |
| 35 | -165 | 1119 | 1974 | 855 | 8 | 86 | INTERFACE_rescored_19.pose.6w41-FabOut.20_0138_0001 |
| 36 | -164 | 1043 | 2237 | 1194 | 5 | 105 | INTERFACE_rescored_11.pose.6w41-FabOut.20_0897_0001 |
| 37 | -161 | 1144 | 2157 | 1012 | 10 | 85 | INTERFACE_rescored_18.pose.6w41-FabOut.19_0255_0001 |
| 38 | -159 | 1130 | 1932 | 801 | 8 | 84 | INTERFACE_rescored_17.pose.6w41-FabOut.24_0100_0001 |
| 39 | -159 | 1054 | 2064 | 1010 | 12 | 85 | INTERFACE_rescored_20.pose.6w41-FabOut.14_0421_0001 |
| 40 | -155 | 1028 | 1933 | 905 | 10 | 78 | INTERFACE_rescored_07.pose.6w41-FabOut.19_0010_0001 |
| 41 | -151 | 1113 | 2002 | 889 | 6 | 82 | INTERFACE_rescored_09.pose.6w41-FabOut.06_0158_0001 |

**SASA: solvent accessible surface area of the mAb-RBD interface**
**SASAh: hydrophobic SASA**
**SASAp: polar SASA**
**nHB: number of hydrogen bonds at the interface**
**nres: number of residues participating in the interface**
